# A rapid genome-wide analysis of isolated giant viruses only using MinION sequencing

**DOI:** 10.1101/2023.03.14.532522

**Authors:** Hiroyuki Hikida, Yusuke Okazaki, Ruixuan Zhang, Thi Tuyen Nguyen, Hiroyuki Ogata

## Abstract

Following the discovery of Acanthamoeba polyphaga mimivirus, diverse giant viruses have been isolated. However, only a small fraction of these isolates has been completely sequenced, limiting our understanding of the genomic diversity of giant viruses. MinION is a portable and low-cost long-read sequencer that can be readily used in a laboratory. Although MinION provides highly error-prone reads that require correction through additional short-read sequencing, recent studies assembled high-quality microbial genomes only using MinION sequencing. Here, we evaluated the accuracy of MinION-only genome assemblies for giant viruses by re-sequencing a prototype marseillevirus. Assembled genomes presented over 99.98% identity to the reference genome with a few gaps, demonstrating a high accuracy of the MinION-only assembly. As a proof of concept, we *de novo* assembled five newly isolated viruses. Average nucleotide identities to their closest known relatives suggest that the isolates represent new species of marseillevirus, pithovirus, and mimivirus. Assembly of subsampled reads demonstrated that their taxonomy and genomic composition could be analyzed at the 50× sequencing coverage. We also identified a pithovirus gene whose homologues were detected only in metagenome-derived relatives. Collectively, we propose that MinION-only assembly is an effective approach to rapidly perform a genome-wide analysis of isolated giant viruses.

## Introduction

Giant viruses are characterized by their remarkably large particles and genomes, some of which overwhelm small unicellular organisms. (La Scola *et al*., 2003; Raoult *et al*., 2004; Philippe *et al*., 2013; Legendre *et al*., 2014). Their large genomes contain genes typically involved in core cellular functions, such as aminoacyl-tRNA synthetases, histones, and fermentation-related genes, thus making them unique among viruses (Raoult *et al*., 2004; Boyer *et al*., 2009; Schvarcz and Steward, 2018; Yoshikawa *et al*., 2019). Following the discovery of Acanthamoeba polyphaga mimivirus, numerous giant viruses have been isolated with diverse morphology, genome size, and genetic content (Aherfi *et al*., 2016; Fischer, 2016; Schulz *et al*., 2022). These giant viruses are classified in the phylum *Nucleocytoviricota* (Koonin and Yutin, 2019; Aylward *et al*., 2021). Recent metagenomic studies have identified uncultured giant viruses from different environments, which revealed the vast diversity and ubiquitous distribution of giant viruses (Endo *et al*., 2020; Moniruzzaman *et al*., 2020; Schulz, Roux, *et al*., 2020). Phylogenetic studies have also revealed numerous gene transfer events from viruses in *Nucleocytoviricota* to cellular organisms (Guglielmini *et al*., 2019, 2022; Irwin *et al*., 2021). These studies highlighted the importance of giant viruses in microbial ecology and evolution.

Metagenomic analysis has revealed the vast diversity of giant viruses, but some isolated viruses are still not represented in the metagenomic data (Schulz, Andreani, *et al*., 2020). Therefore, virus isolation is an indispensable approach to characterize the diversity of these viruses. Most giant viruses have been isolated with a co-culture method by using free-living amoebae, such as *Acanthamoeba* and *Vermamoeba* species, as a host. This method is well established and has been used to isolate many viruses (La Scola *et al*., 2010; Abergel *et al*., 2015; Boudjemaa *et al*., 2020). These viruses were characterized and classified according to their morphological features observed by electron microscopy, particle and DNA content profiles analyzed by flow cytometry, and molecular phylogeny based on conserved genes (Boughalmi *et al*., 2013; Khalil *et al*., 2016; Aoki *et al*., 2019; Sahmi-Bounsiar *et al*., 2021). Although electron microscopy and flow cytometry can identify new giant viruses with atypical morphological or physical properties, these methods are unable to distinguish closely related viruses with similar morphology and genome sizes. Molecular phylogeny of specific genes can resolve the phylogenetic relationships of giant viruses with their close relatives, but this approach does not reveal their genomic features (Yutin *et al*., 2013; Aherfi *et al*., 2018). Genome sequencing analysis is therefore essential to characterize newly isolated giant viruses on the basis of functional repertoire and taxonomy. Whole genome data are, however, available only for a small fraction of isolated viruses because of the time and cost involved for whole-genome sequencing.

MinION (Oxford Nanopore Technologies, Oxford UK) is a portable and low-cost long-read sequencer that can be readily implemented in a laboratory. Its installation cost currently starts from 1,000 USD, which is much lower than that of other high-throughput sequencing platforms. Long reads of nanopore sequencing can elucidate repeat structures that cannot be resolved by short-read sequencing and produce longer contigs than that achieved with the short-read-only assembly. However, the long reads are noisy with numerous base-calling errors, which usually require correction with highly accurate short reads. Long- and short-read sequences mutually compensate for the disadvantages of each technology, and some giant viruses have been sequenced by the hybrid assembly of both sequencing platforms (Yoshida *et al*., 2021; Xia *et al*., 2022). Recently, however, several studies have demonstrated that the nanopore long-read sequence can produce highly accurate genomes for bacteria and yeast without the requirement for short-read correction (Loman *et al*., 2015; Istace *et al*., 2017). The bacteria genomes showed 99.5% nucleotide identity to the reference genome, while the yeast genome showed 99.8% identity to the reference genes. These results imply that the MinION allows whole-genome sequencing for giant viruses more rapidly at a lower cost than hybrid assembly. However, the effectiveness of the long-read-only sequencing approach for giant viruses has not yet been assessed.

In the present study, we evaluated the quality of genomes assembled only by MinION sequencing. The accuracy of the sequencing method was assessed by re-sequencing a prototype of the family *Marseilleviridae*, marseillevirus marseillevirus (MsV) T19. The genome was assembled with four assemblers at different coverages, and the performance of these assemblers was compared. As a proof of concept, we assembled genomes of five newly isolated viruses using only MinION sequencing and identified them as two marseilleviruses, one pithovirus and two mimiviruses. Their average nucleotide identity (ANI) to the known isolated viruses suggests that the newly isolated viruses represent new species. Collectively, the present study demonstrated that the long-read-only sequencing approach is a rapid and low-cost option to explore the genomic diversity of giant viruses.

## Experimental procedures

### Cells and viruses

*Acanthamoeba castellanii* (Douglas) Page, strain Neff (ATCC 30010) was maintained with peptone-yeast extract-glucose (PYG) medium at 28°C and used as a host for giant viruses. MsV T19 was used as a marseillevirus prototype (Boyer *et al*., 2009).

### Sample collection

Sediment and water samples were collected from Lake Biwa, Japan, on July 16th, 2021. Sediments were collected from the bottom of the north (35.13.2152 N, 135.59.7862 E) and south (35.00.4823 N 135.54.0953 E) basins using a core sampler. The collected cores were approximately 400 mm in height and were divided into three layers: top, middle, and bottom. Each sediment sample was resuspended in Page’s amoeba saline (PAS). Large particles were removed from the sample by filtration using a Whatman No. 43 filter paper (GE Healthcare). Water samples were collected at 60 m depth from the north basin and subsequently filtered through a 5-μm filter (Millipore) and a 0.22-μm Sterivex cartridge (Merck). The 0.22-μm filter was then removed from the cartridge and vortexed in approximately 30 mL of PAS until the filter turned white.

### Virus isolation

The water and sediment samples were inoculated into amoeba culture that were seeded onto 96 well plates at the density of 1 × 10^3^ cells per well with PYG medium. Each well contained 200 μL solution of 1×penicillin/streptomycin (Wako), 25 μg/mL of amphotericin B, 100 μg/mL of ampicillin, 20 μg/mL of ciprofloxacin, and 60 μL of an environmental sample. Five wells that showed cytopathic effect were collected and purified by the end-point dilution method. Each culture was identified as an isolated virus and designated as BNT8A, BSD11G, BST12E, BST10G, and BN60m3A.

### Negative staining

Cultured viruses were collected by centrifugation at 9,000 rpm for 1 h at 4°C (Sorvall ST8FR, Thermo Scientific) and resuspended in phosphate-buffered saline. The virus suspension was fixed in 1% glutaraldehyde solution for several hours. The fixed samples were then transferred onto a collodion mesh (Nisshin-EM). The viral particles were stained with uranyl acetate and observed by an H-7650 transmission electron microscope (Hitachi).

### DNA extraction

MsV T19 was cultured in a T25 flask, and the viral particles were collected as described above. The viral particles were treated sequentially with 2.5 mg/mL lysozyme, 2.8 mg/mL bromelain, and 2 mg/mL proteinase K with 1% SDS. Each treatment was performed overnight. DNA was extracted by treatment with phenol for two times and treatment with chloroform for two times, followed by ethanol precipitation.

For newly isolated viruses, different extraction methods were applied depending on their particle morphology. DNA of BNT8A, BSD11G, and BST12E was extracted as described above. BST10G and BN60m3A were cultured in a T75 flask, collected by centrifugation at 9,000 rpm for 15 m at 4°C, and treated with 50 mM NaOH at 95°C for 5 m. 10% of 1 M Tris-HCl at pH 8.0 was added, followed by dilution with TE buffer at pH 8.0. DNA was extracted by treatment with phenol for three times and treatment with chloroform for three times, followed by ethanol precipitation.

### Nanopore sequencing

Genomic DNA concentration was measured by a Qubit 4 fluorometer using Qubit dsDNA HS and BR Assay Kits (Invitrogen). A sequence library was prepared using the Ligation Sequencing Kit (SQK-LSK109, Oxford Nanopore Technologies) from 2.25 μg of MsV genomic DNA, following the manufacturer’s instructions. For the five newly isolated viruses, a multiplexed library was prepared using the Rapid Barcoding Kit (SQK-RBK004, Oxford Nanopore Technologies) from 100 ng of each genomic DNA for BNT8A, BSD11G, and BST12E and 8 ng of each genomic DNA for BST10G and BN60m6A. Each library was sequenced using one MinION flow cell (R9.4.1) (Oxford Nanopore Technologies) on MinION Mk1C with MinKNOW v.21.05.12 software (Oxford Nanopore Technologies).

### Genome assembly

For MsV genome assembly, base-calling was performed by the fast and high-accuracy modes in Guppy (v5.0.12) and assembled using four assemblers with default parameters: Flye (v2.8.2) (Kolmogorov *et al*., 2019), Miniasm (v0.3) (Li, 2016), Raven (v1.5.0) (Vaser and Šikić, 2021), and Wtdbg2 (v2.5) (Ruan and Li, 2020). The assembled contigs were polished using long reads with three rounds of Racon (v1.4.13) (Vaser *et al*., 2017), followed by Medaka (v1.4.1) (https://github.com/nanoporetech/medaka). Raw sequencing data were base-called by the fast and high-accuracy modes and then subsampled by Seqkit (Shen *et al*., 2016) at 0.1%, 0.2%, 0.5%, 1%, 2%, 5%, 10%, and 20%, corresponding to approximately 5×, 10×, 25×, 50×, 100×, 250×, 500×, and 1000× coverage, respectively. The subsampling process was repeated five times with different random seeds. Reads in each subsampling process were assembled independently.

For newly isolated viruses, the demultiplexed raw data were base-called by the high-accuracy mode using Guppy, followed by assembly and polishing as described above. The reads were subsampled at 1%, 2%, 5%, 10%, 20%, and 50% of the total reads and assembled using Flye, Miniasm, and Raven followed by polishing as described above.

### Quality assessment of MsV assembly

As MsV assemblies included putative fragmented contigs, a reciprocal comparison between all the contig pairs was performed in each assembly by using BLASTN (v2.0.11) (Camacho *et al*., 2009) with the word size option set as 100. In each pair, a smaller contig showing 99% identity to the larger one was excluded from the dataset. The resulting nonredundant contigs were compared with the T19 reference sequence (GenBank: GU071086.1) by using BLASTN. Sequence identity was determined as the number of identical bases divided by the alignment length. Genome fraction was defined as the proportion of the reference genome covered by the alignments. Gap length was calculated as the sum of the gap length that appeared in all alignments. The alignments were visualized using custom Python scripts.

The accuracy of the predicted protein-coding sequences was assessed as follows. Protein-coding genes were predicted from each assembly using Prodigal (v 2.6.3) (Hyatt *et al*., 2010). To eliminate the effect of gene prediction methods, protein-coding genes were predicted *de novo* from the reference MsV T19 genome. When an amino acid sequence exhibited the same length and 100% sequence identity to a gene predicted in the reference genome, the protein-coding sequence was assumed to be precisely predicted.

### Characterization of putative repeat regions found in MsV assembly

Raw long reads generated by the high-accuracy base-calling mode were mapped to the T19 reference sequence using Minimap2 (v2.22) (Li, 2016). The mapping profile was converted to the BAM format by using SAMtools (v1.15) (Li *et al*., 2009). The resultant BAM file was converted to the WIG format and visualized using the Integrative Genome Viewer (v2.13.2). Putative repeat regions reported in a previous study (Bryson *et al*., 2022) were manually determined to be located in the nucleotide positions from 18,816 to 35,332 and from 317,602 to 319,352. Among the raw reads that were mapped to the repeat regions, those longer than the length of the regions were identified. The number of repeat units included in these reads was determined by MUMmer (v4.0.0) (Marçais *et al*., 2018) by using the repeat unit as a query.

### Comparison to known viruses

The ANI between genomes of known and newly isolated viruses was calculated using FastANI (v1.3.3) (Jain *et al*., 2018). Clustering and visualization were performed using custom Python scripts. Genomes of the analyzed giant viruses were retrieved from the NCBI Virus database (Table S1).

### Comparison to hybrid assembly

Genomic DNA of the newly isolated viruses was extracted as described above and sequenced using the Illumina NovaSeq 6000 system with the 150-bp paired-end mode. A total of 1 GB data were obtained for each virus. The quality control of genomic DNA, library preparation, and sequencing were performed by Rhelixa, Inc. (Japan). The Medaka-polished genomes were further polished using Pilon (v1.23) (Walker *et al*., 2014) with short reads, and protein-coding genes were predicted as described above. A BLASTP search was performed using the predicted genes with the Pilon-polished genomes as queries and genes in the long-read-only assembly as subjects. The presence, identity, and coverage of the genes in the long-read-only assembly were investigated as follows: (1) both query coverage and identity were 100% – the gene was assumed as “complete”; (2) the query coverage was < 100% but the identity was 100% – the gene was defined as “truncated”; (3) the identity was < 100% but the query coverage was 100% – the gene was classified as “mismatch”; (4) queries had a BLAST hit but were not specific for the above case – the gene was classified as “presence”; and (5) queries had no BLAST hit – the gene was determined as “absence.”

### Comparison of genomes between assemblers

Genomes of the newly isolated viruses assembled by Flye, Miniasm, and Raven were compared with each other by using MUMmer (v4.0.0). Short contigs found in the BST12E assemblies reconstructed by Raven and Miniasm were characterized using BLASTX (v.2.13.0) against the NR database and excluded from the analysis as contaminants of host DNA.

### Genomic analysis of the newly isolated pithovirus BST12E

To identify genes specific to pithovirus BST12E, orthologous genes among the three isolated pithoviruses, including BST12E, were detected using OrthoFinder (v2.5.2) (Emms and Kelly, 2019). Protein sequences of pithovirus sibericum was retrieved from NCBI (GenBank Accession No.: KF740664.1). Protein sequences of pithovirus massiliensis were predicted from a genome sequence (GenBank Accession No.: LT598836.1) by Prodigal, as its protein sequences were unavailable in GenBank.

Genes specific to BST12E were annotated using DIAMOND BLASTP (v2.0.11) (Buchfink *et al*., 2021) against the NR database. InterProScan (Jones *et al*., 2014) was used to search known domains or motifs in two BST12E-specific proteins BST12E_145 and BST12E_501. Their homologous proteins were retrieved through a BLASTP search against the NR database. The homologous sequences were aligned using MAFFT (v7.487) (Katoh and Standley, 2013), followed by the construction of phylogenetic trees for the two proteins with LG+I+G4 and Blosum62+I+G4 model, respectively, using IQ-TREE (v2.1.3) with ModelFinder (Kalyaanamoorthy *et al*., 2017; Minh *et al*., 2020). The phylogenetic trees were visualized by ITOL (v6) (Letunic and Bork, 2021).

## Results

### Assembly performance for the prototype marseillevirus only using long-read sequencing

Genomic DNA of MsV T19 was sequenced with MinION, by using the Ligation Sequencing Kit (Table. S2). Base-calling by the fast and high-accuracy modes generated 2.03 and 1.99 GB sequence reads from one run of the MinION flow cell, respectively. The data were subsampled and assembled independently five times with the four assemblers at different coverages. Although each assembly showed varied accuracy, all assemblers recovered at least one nearly complete assembly that covered 99.9% of the reference genome with >99.9% identity at 50× or higher coverages (Fig. 1 and Supplementary data). Among the four assemblers, Flye showed stable performance. All assemblies by Flye at 50× or higher coverages covered more than 99.99% of the reference genome with >99.98% identity without polishing (Supplementary data). At the lower coverages, Flye assembled longer contigs, while Miniasm assembly showed higher identity than those of Raven and Wtdbg2. Wtdbg2 assembly was less accurate, particularly at higher coverages, compared to the other assemblers.

**Figure 1.**
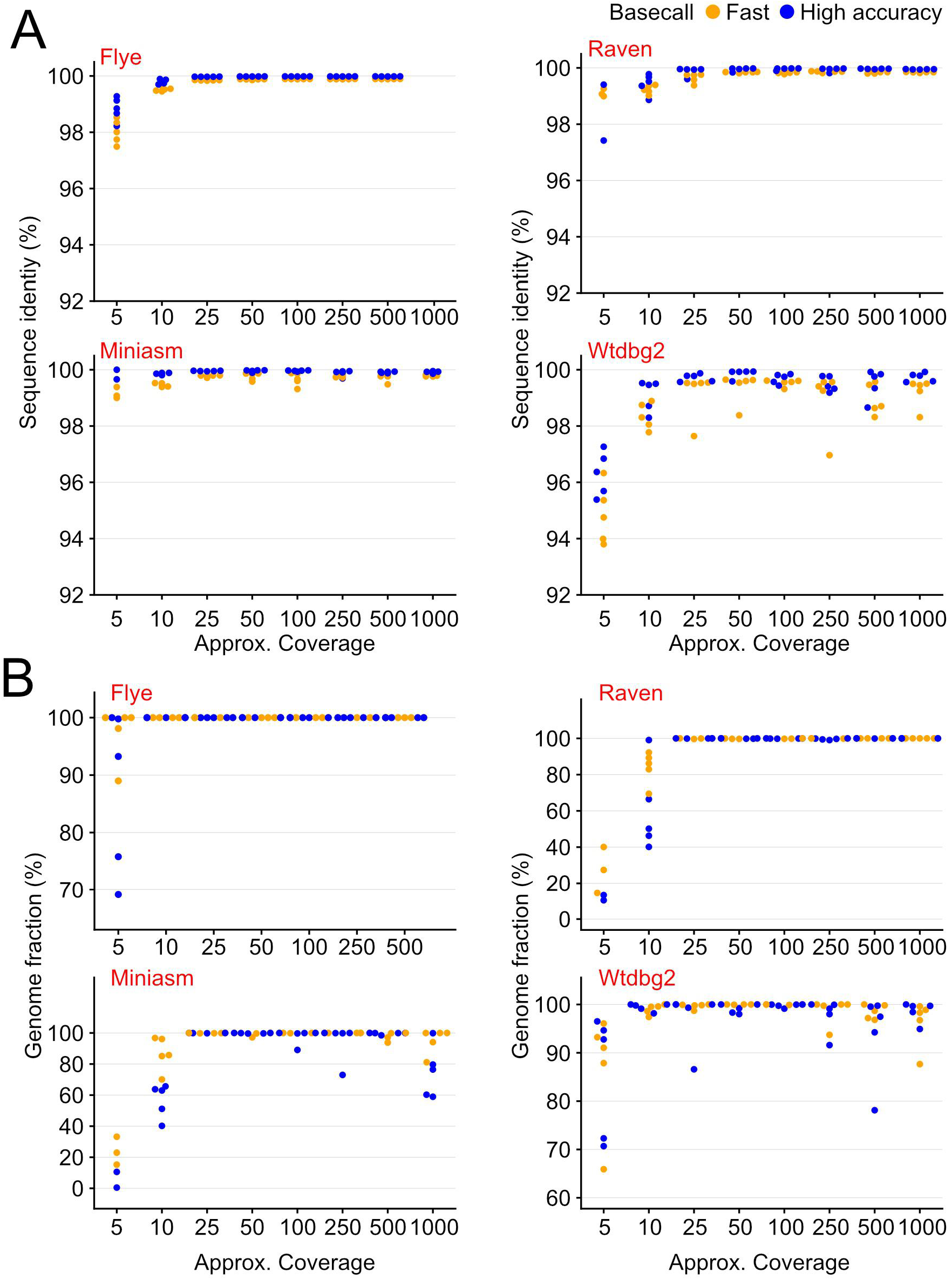
(A) Sequence identity between marseillevirus marseillevirus (MsV) assemblies at different coverages and the reference genome. (B) Fraction of the reference genome covered by genomes assembled at different coverages. (A, B) Names of software are shown in red. Dots indicate each assembly. Orange and blue dots indicate fast and high-accuracy base-call modes, respectively. Subsampling was performed five times at each coverage with different random seeds.

Some assemblies were longer than the reference genome (Fig. S1). The MsV genome is circular, but only Flye implements circularization of assembled genomes. Therefore, some genomes assembled by Raven, Miniasm, and Wtdbg2 harbored an overlapping region at their termini, which affected the assembly size (Fig. S2). Furthermore, some assemblies contained tandem repeats of approximately 16 and 1.7 kbp (Fig. S2). These repeats were located in the regions that were previously identified as putative repeat regions by short-read mapping (Bryson *et al*., 2022). This previous study indicated that these regions were expanded during successive passages. To further investigate these repeats, we counted the number of repeat units included in the raw long reads. The raw reads mapped to the repeat regions contained a different number of repeat units (Fig S3). These results suggest that the sequenced virus had variation in the copy number of repeat units, resulting in longer assemblies.

Although highly accurate assemblies were reconstructed at 50× coverage, 100× coverage resulted in further improvement. The number of gaps decreased, and the number of precisely predicted proteins increased at 100× coverage, compared to those at 50× coverage (Fig. 2). At 100× coverage, Medaka polishing resulted in less gaps and a higher number of precisely predicted proteins than genomes polished by Racon or those not subjected to polishing (Fig. 2). The quality reached a plateau at around 100× coverage regardless of polishing (Fig. S4). Although no genome was identical to the reference genome, these results indicate that higher coverage and polishing improves the quality of genomes until around 100× coverage.

**Figure 2.**
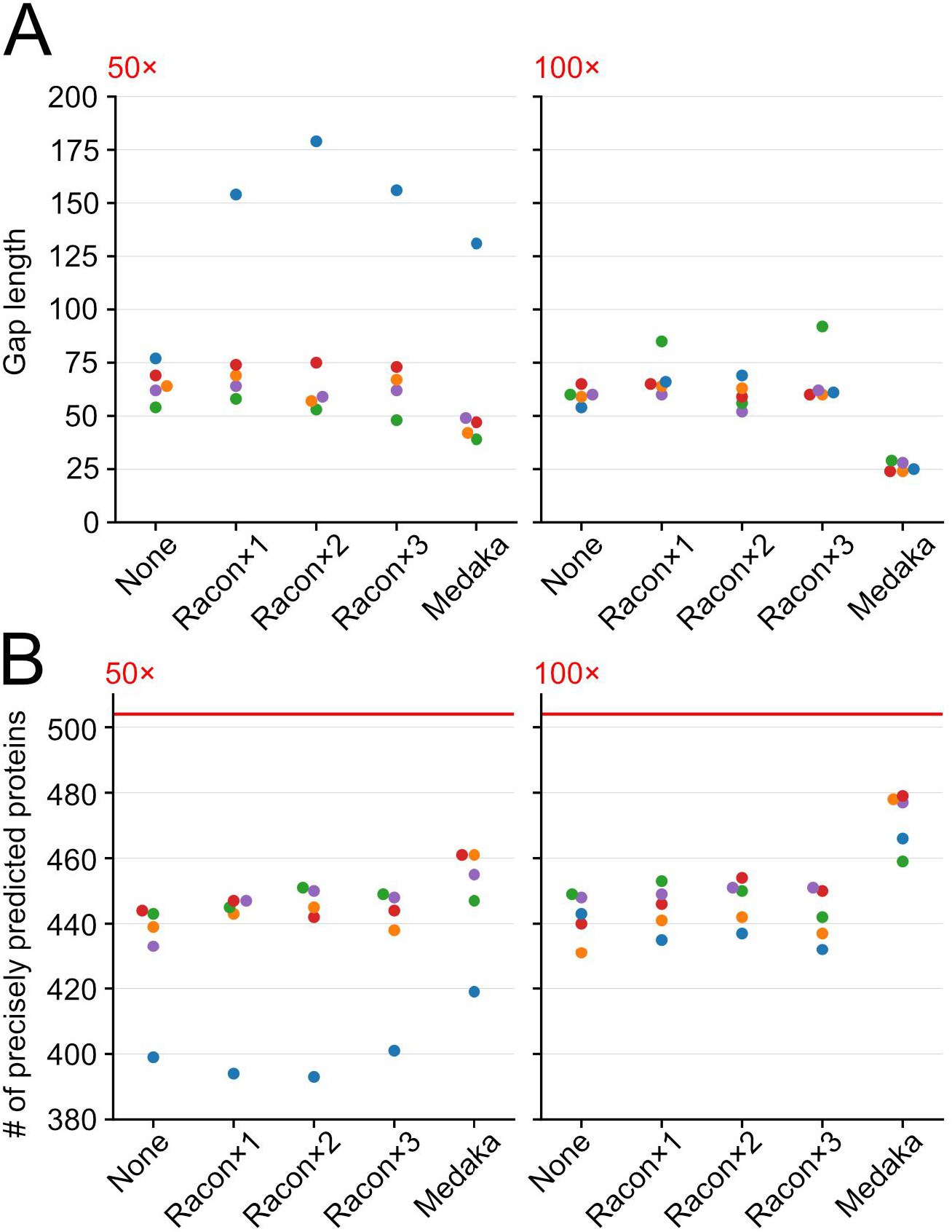
Comparison of the quality of MsV genomes derived by Flye assembly with or without polishing at 50× and 100× coverages. (A) Gap length. (B) The number of precisely predicted proteins as defined in Materials and Methods. Red lines indicate the number of genes predicted in the reference genome. X-axes indicate the method of polishing. Colors correspond to each subsampling process with different random seeds.

### Morphological characterization of giant viruses isolated from Lake Biwa

Five viruses were isolated from different sources collected at Lake Biwa, Japan (Table. S3).

Based on the results of negative staining, these viruses were morphologically classified into three types (Fig. 3). The first type (BNT8A and BSD11G) exhibited a polyhedron shape with approximately 300 nm diameter, similar to marseilleviruses. The second type (BST10G and BN60m3A) was larger than 500 nm in diameter and had fibril-like structures, similar to mimiviruses. The third type was BST12E with a large oval-shaped virion of over 1 μm in length.

**Figure 3.**
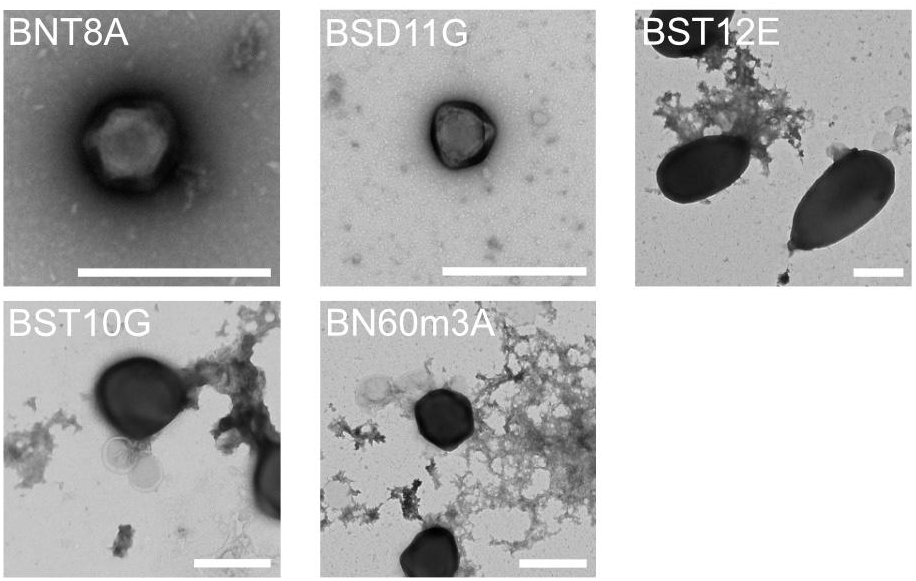
Negative staining images of newly isolated viruses from Lake Biwa. Bars indicate 500 nm. Isolate names are shown on the top left.

### Long-read-only assemblies of the isolated viruses

The isolated viruses were sequenced using the Rapid Barcoding Kit. This kit enables fast library preparation and multiplexed sequencing, thereby reducing the time and cost of sequencing compared to the Ligation Kit. The N50 of the sequence reads was shorter than that of MsV T19, which may be because of the difference in library preparation (Table S2). Nevertheless, a single circular contig was obtained for three viruses, namely BNT8A, BSD11G, and BST12E (Table S4). BNT8A and BSD11G were assembled into a single contig by all assemblers. BST12E was split into long and short contigs by Raven and Miniasm. These short contigs seemed to be a fragment of host mitochondria based on the BLASTX search (Table S5). BST10G and BN60m3A were split into several contigs in all assemblies. Among the contigs of these two mimivirus-like viruses, only one contig of BN60m3A assembled by Raven have length (1.14 Mb) comparable to the typical mimivirus genome size.

### Quality of long-read-only assemblies of the isolated viruses

We investigated the completeness of protein prediction in long-read-only assembled genomes by comparing the predicted proteins with those in the genomes polished by short-read sequencing (Fig. 4 and Table S6). Nearly 90% of the genes in the hybrid assemblies were recovered with 100% identity and coverage in the long-read-only assemblies. Only <2% of the genes in the hybrid assembly were absent in long-read-only assemblies. These results indicate that the long-read-only assemblies using multiplexed sequencing can provide an overview of gene content with a high-accuracy.

**Figure 4.**
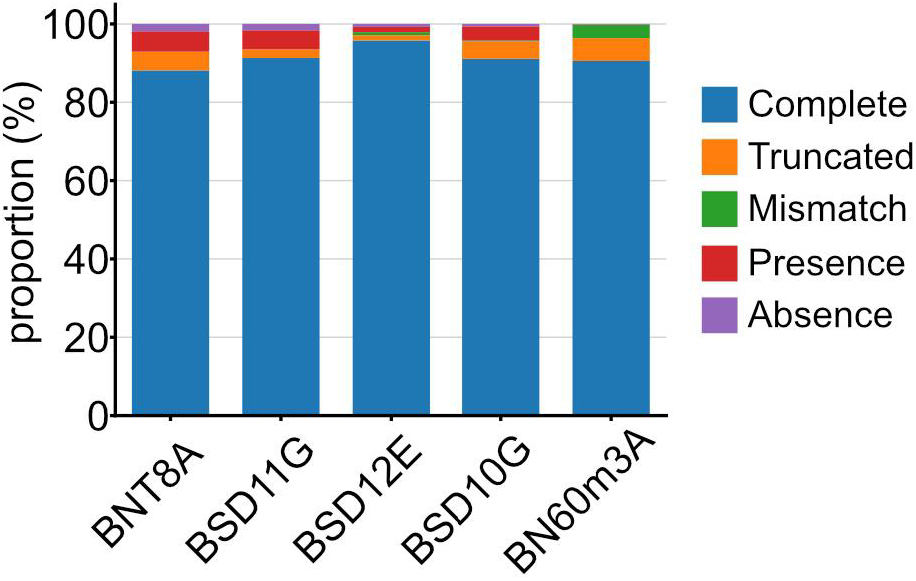
Accuracy of protein prediction in the long-read-only assembly of newly isolated viruses. Each bar shows the proportion of proteins in hybrid assembly genomes that were classified according to their prediction accuracy defined in Materials and Methods. Coverage of assemblies are 501×, 495×, 439×, 237×, and 223× for BNT8A, BSD11G, BST12E, BST10G, and BN60m3A, respectively. Counts are shown in Table S6.

To investigate consistency between the assemblers, assemblies obtained by Flye, Miniasm, and Raven, were compared with each other (Figs. S5 and S6). Wtdbg2 was excluded because of its unstable performance in MsV assembly. The assemblies of BNT8A and BSD11G were highly consistent between the assemblers and polishing with short reads further improved the consistency up to 100% in some comparisons. Other assemblies (BST12E, BST10G, and BN60m3A) also showed high nucleotide identity >99.9% after polishing but never reached 100%. Moreover, in several comparisons, a few percent of genomes were not aligned with other assemblies, thus indicating that some assemblers missed some portions of the genomes.

Genomes of the newly isolated viruses were also assembled at different coverages by using Flye, Miniasm, and Raven. For BNT8A, BSD11G, and BST12E, approximately 20×-to 30×-coverage data recovered most parts of the assemblies that were reconstructed using the original read population (Fig. S7). These genomes recovered most of the proteins in the hybrid assemblies, over 50% of which were completely recovered (Fig. 5). At 50× coverage, Flye and Raven recovered entire genomes with continuous contigs, where 90% of proteins were completely recovered. Genomes of BST10G and BN60m3A were recovered at 50× coverage, but they were more fragmented than those of BNT8A, BSD11G, and BST12E (Fig. S7). Nevertheless, most of the proteins were recovered, and >50% of them were complete (Fig. 5). Collectively, our results indicate that the genomic compositions of these giant viruses can be analyzed at 50× coverage.

**Figure 5.**
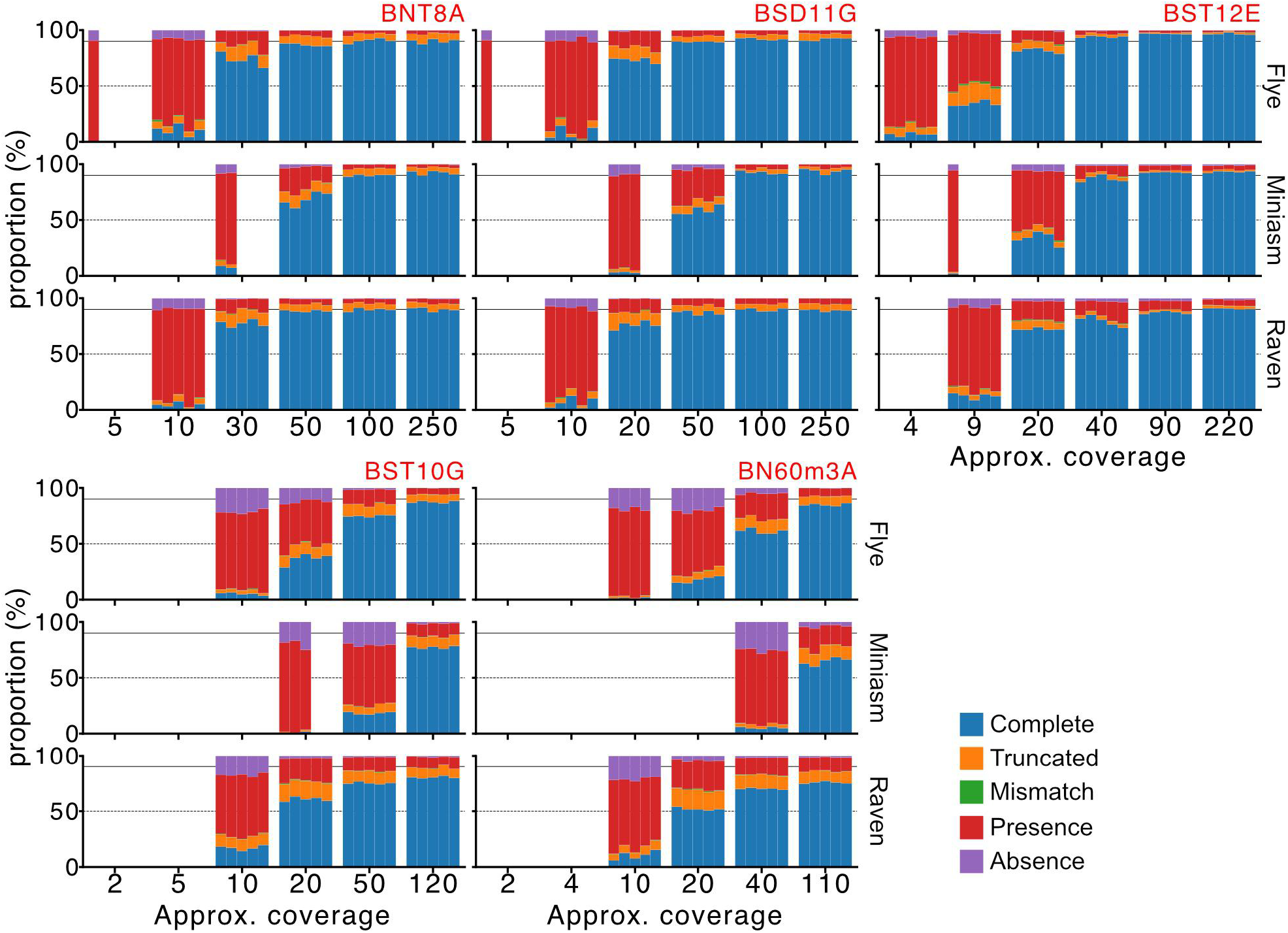
Accuracy of protein prediction in genomes of newly isolated viruses assembled at different coverages by using Flye, Miniasm, and Raven. Protein-coding genes in long-read-only assemblies were compared with those in hybrid assembly. Each bar indicates an assembly. At some assemblies, no contig was reconstructed. X-axes show approximate coverages for each assembly. Each row represents the assembler, namely Flye, Miniasm, and Raven, from top to bottom. Solid and dashed horizontal lines indicate 90% and 50%, respectively. Red labels show the virus isolate names.

### Genome-based taxonomic classification of the isolated viruses

To taxonomically classify the newly isolated viruses, a long-read-only assembly of each virus was compared with genomes of isolated giant viruses deposited in the NCBI Virus database. As Flye showed the most stable performance for MsV T19 and all the newly isolated viruses, we used the assembly by Flye for the comparison. The ANI was consistent with the morphological observation (Table 1 and Fig. S8). BNT8A and BSD11G showed the highest ANI to marseilleviruses. Similarly, the closest relative of BST10G and BN60m3A was a mimivirus. The two newly isolated mimiviruses were closely related, and BST10G showed 99.22% ANI to BN60m3A (Fig. S8). The large oval-shaped virus BST12E showed similarity with pithovirus sibericum, a prototype of pithovirus. In bacteria, 95% ANI is considered as a species boundary (Jain *et al*., 2018). This criterion was also used in recent taxonomic classification of giant viruses, as this ANI value is considered a useful metric to classify viruses (Bobay and Ochman, 2018; Aylward *et al*., 2021). All the newly isolated viruses in the present study showed ANI below 95% to the reference sequences, suggesting that they represented new species. The hybrid assemblies also confirmed the taxonomic classification and novelty of the isolated viruses (Table S7). We tentatively named the viruses as shown in Table S3.

ANI was also compared for genomes assembled with subsampled reads by using Flye, Miniasm, and Raven (Fig. 6). The ANI values were constant for the genomes assembled at different coverages or by different assemblers. In particular, the values were almost the same at around 50× or higher coverages. The closest viruses varied between the assemblies as some reference viruses showed extremely high ANI with each other (Fig. S8). Collectively, these results indicate that a coverage of 50× is adequate to resolve the classification of giant viruses at the species level.

**Figure 6.**
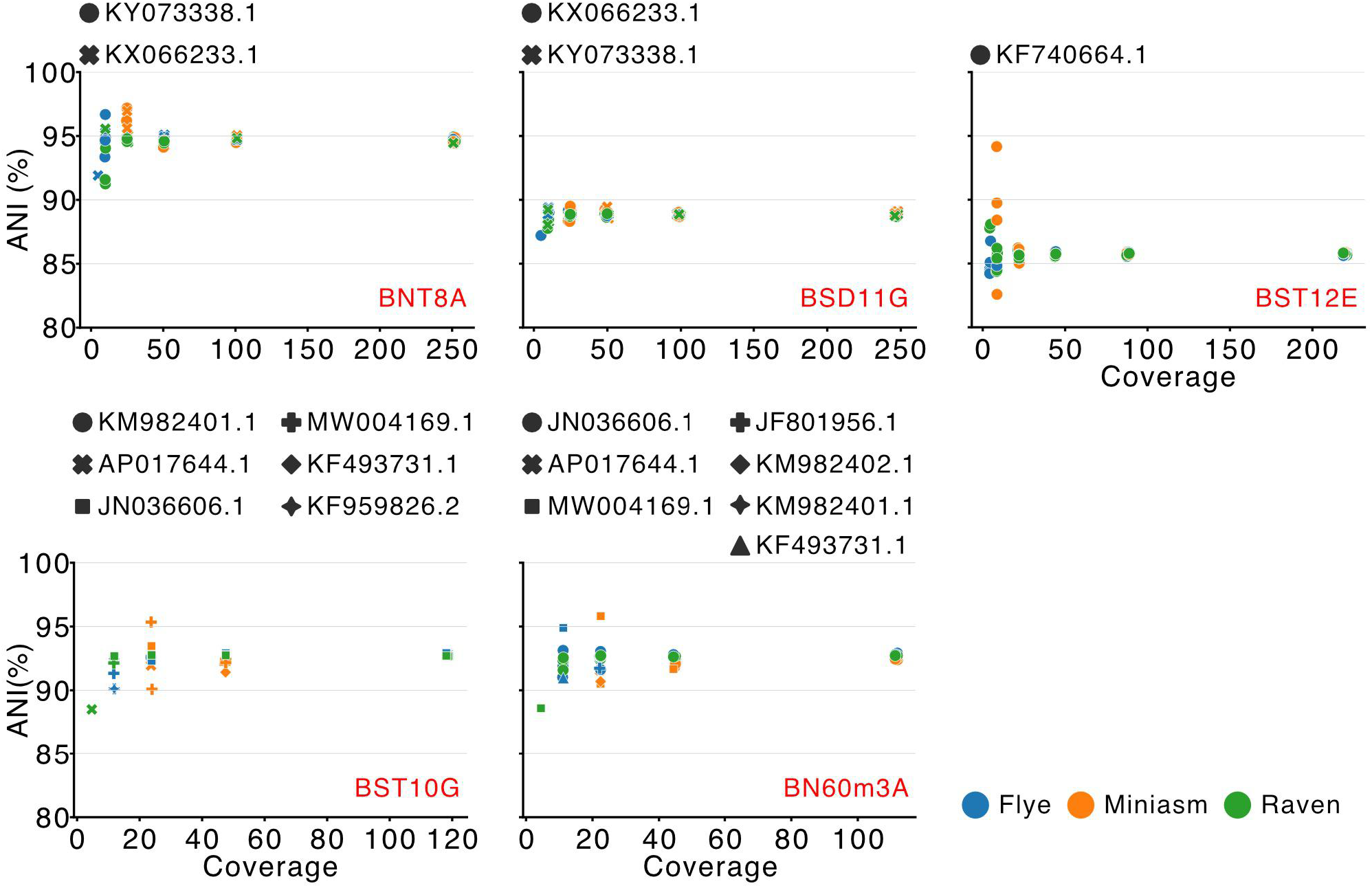
ANI values of the genomes of newly isolated viruses assembled at different coverages by using Flye, Raven, and Miniasm, to the closest reference viral genomes. Dots, colors, and shapes indicate subsampling, software, and reference viruses that showed the highest ANI value, respectively. Red labels show the isolate names for the newly isolated viruses.

### Analysis of genes specific to the pithovirus BST12E

Currently, three genomes of pithoviruses are available including BST12E (Legendre *et al*., 2014; Levasseur *et al*., 2016). BST12E showed 85.7% and 90.2% ANI to the reference pithovirus sibericum and pithovirus massiliensis, which is not yet included in the NCBI Virus database, respectively (Table S8). The low ANI to the known viruses suggests that BST12E has distinct genomic features. Therefore, we investigated the long-read-only assembly of BST12E reconstructed by Flye. We identified 20 of 525 BST12E genes that are specific to BST12E (Table S9). The BLASTP search against the NCBI NR database revealed that 18 of the BST12E-specific genes were ORFans without any homologue. The remaining two genes, namely BST12E_145 and BST12E_501, were homologous to those of bacteria and marseillevirus, respectively. InterProScan identified that BST12E_145 belongs to the clavaminate synthase-like superfamily. According to the InterProScan results, BST12E_501 did not show any motif, although it showed homology to a marseillevirus R3H domain-containing protein (Table S9). Protein homologous to these two proteins were retrieved from the NCBI NR database, and their phylogenetic relationship was investigated (Fig. S9). In addition to bacteria, BST12E_145 homologues were encoded in metagenome-derived pithoviruses detected in sediments from the Loki’s Castle hydrothermal vent area (Bäckström *et al*., 2019). A homologue of BST12E_501 was also encoded in the genome of Orpheovirus. Orpheovirus is a member of the order *Pimascovirales*, which includes pithoviruses and marseilleviruses (Aylward *et al*., 2021). Taken together, we demonstrated that our method can reveal distinct genomic features of isolated giant viruses.

## Discussion

MinION is a low-cost and portable sequencing platform that can be easily installed in a laboratory and the cost of its installation is low compared to that of other high-throughput sequencing platforms. The present study evaluated the performance of MinION sequencing for genomic characterization of isolated giant viruses. Re-sequencing of a prototype marseillevirus revealed that some assemblers can reconstruct high-quality genomes with 99.98% identity and over 99.99% coverage at 50× or higher coverages (Fig. 1). Polishing of the assembled genomes by using original long reads further improved the quality by reducing the number of gaps and increasing the number of precisely predicted proteins (Fig. 2). The number of gaps was reduced at 100× coverage compared to that at 50× coverage, indicating that higher coverages enabled more accurate polishing (Figs. 2 and S4). Overall, we conclude that MinION sequencing reconstructs highly accurate genomes of a giant virus without the requirement for correction with short reads.

As a proof of concept, we applied the long-read-only assembly for *de novo* sequencing of newly isolated viruses. Consistent with morphological observations, two marseilleviruses, one pithovirus, and two mimiviruses were identified on the basis of ANI values (Figs. 3 and S8 and Table 1). Approximately 90% of the proteins were precisely predicted in long-read-only genomes, indicating that gene composition analysis is possible with the long-read-only assembly (Fig. 4). Assemblies of subsampled reads demonstrated that most of the genes and genomes were recovered at approximately 50× coverage, allowing to perform genome-wide analysis and taxonomic classification based on ANI (Figs. 5, 6, and S7). The maximum giant virus genome reported to date is 2.5 Mbp of pandoraviruses (Philippe *et al*., 2013), suggesting that around 100 MB data allow analysis of taxonomy and genomic composition for one giant virus. Our results demonstrated that multiplexed sequencing easily met this data size for five viral isolates using a single MinION flow cell (Table S2). This data size would also be adequate for the Flongle flow cell, a down-scaled version of the MinION flow cell, which costs 90 USD and generates sequences of up to 2.8 GB data. Furthermore, a newer version of the MinION flow cell (R10.4) was recently commercialized. This flow cell generates more accurate reads (Sereika *et al*., 2022), which improves the assembly quality and reduces size of the required data. These technologies may further reduce the cost of long-read-only genome-wide analysis of giant viruses. Collectively, our study demonstrated that nanopore sequencing is a rapid and highly cost-effective approach to explore the genomic diversity of giant viruses.

The genome quality of long-read-only assemblies was not perfect and should be carefully examined before their deposition in the public databases. We showed that even at a high coverage, no genome was identical to the reference, and a few gaps remained (Fig. S4). This result is consistent with the findings of previous studies assembling microbial genomes using only nanopore sequencing (Loman *et al*., 2015; Istace *et al*., 2017). Some assemblers also provided genomes longer than the reference genome (Fig. S1). Longer genomes were partially explained by the lack of a circularizing step in some assemblers, leading to overlapped regions at the terminals (Fig. S2). We also found that a sequenced virus had repeat regions which may have carried a varying number of repeat units (Fig. S3). The regions were previously predicted by short-read assembly and considered as the region expanded during successive passages (Bryson *et al*., 2022). These results showed that long-read sequencing is advantageous to resolve repeat regions, but they also suggest that some software may be affected by minor variants. Furthermore, *de novo* assembly of isolated viruses showed slight differences between the software used, particularly in the aligned proportion that was not corrected by short-read polishing (Figs. S5 and S6). This result suggests that some regions were missing or redundant in some assemblies.

The newly isolated viruses showed an ANI of <95% to the known viruses, suggesting that they represent new species (Table 1 and S7). Among them, the newly isolated pithovirus BST12E was distinct from the other isolated pithoviruses based on ANI and encodes 20 genes whose homologs were absent in the other isolated pithoviruses (Table S9). Interestingly, two of these genes were detected in the genomes of some other members of the order *Pimascovirales*, including metagenome-derived sequences (Fig. S9). This result suggests that ancestral viruses in *Pimascovirales* had these genes, which were lost in the other isolated pithoviruses. Alternatively, the genes may be transferred to BST12E by other microorganisms including viruses. These results indicate that the long-read-only assemblies could highlight the diversity and evolutionary history of giant virus genomes.

During the two decades after the first mimivirus was reported, new giant viruses were constantly isolated with remarkable morphological and genomic features (Fischer, 2016; Abrahão *et al*., 2018; Yoshikawa *et al*., 2019; Boratto *et al*., 2020). In contrast, genome-wide data are limited for viruses showing similar morphology and/or marker genes to previously sequenced viruses. The present study proposed a cost- and time-effective pipeline for genome-wide analysis of giant viruses by MinION sequencing. Application of this approach revealed that viruses phylogenetically close to previously characterized viruses still have distinct genomic features. A recent metagenomic analysis revealed that although giant viruses are ubiquitous in the ocean, their distribution exhibits a heterogeneous pattern (Endo *et al*., 2020). Here, we isolated new viruses from a freshwater lake. Previous studies have isolated giant viruses from terrestrial environments (Yoosuf *et al*., 2014; Schulz, Andreani, *et al*., 2020). These environments are spatially more heterogeneous than the ocean and may encompass distinct diversity of giant viruses. Increased sampling efforts and genomic analysis of giant viruses isolated from various environments may highlight their genomic diversity, thereby revealing ecological roles and evolutionary histories of giant viruses.

## Acknowledgement

We thank Dr. Bernard La Scola for kindly providing us marseillevirus marseillevirus T19. Field sampling was supported by Center for Ecological Research, Kyoto University, a Joint Usage / Research Center. The electron microscopy study was supported by Division of Electron Microscopic Study, Center for Anatomical Studies, Graduate School of Medicine, Kyoto University. Computation time was provided by the SuperComputer System, Institute for Chemical Research, Kyoto University.

## Data Availability Statement

The raw sequence data are available at DDBJ with accession number DRA015450. The daft genome sequences for the isolated viruses are available at The GenomeNet FTP site (ftp://ftp.genome.jp/pub//db/community/GV_draft_genomes/Hikida_et_al_2023).

## Funding Statement

This study was supported by Japan Society for the Promotion of Science KAKENHI grant number 22K15175, 21J00174 to HH, 22H00384, 19H05667, 18H02279 to HO.

## Conflict of Interest Disclosure

The authors declared no conflict of interest.

## Author Contribution

HH and YO conceptualized the study. HH, YO, RZ, and TTN performed investigation. HH and HO acquired financial support. HH performed formal analysis. HH, YO, and OH write original draft. All authors reviewed, edited, and finalized the draft.

## Figure legends

**Table 1**.

Giant virus genomes showing the highest ANI value with long-read-only assembly of newly isolated viruses.

## Supporting information

**Figure S1**

Length of MsV genomes assembled at different coverages. Red lines indicate the size of the reference MsV T19 genome. Names of software are shown in red. Dots indicate each assembly. Orange and blue dots indicate fast and high-accuracy base-call modes, respectively. Subsampling was performed five times at each coverage with different random seeds.

**Figure S2**

Genome-wide comparison between genomes assembled in this study at 100× coverage and the reference genome, which are shown in the X-axis and Y-axis, respectively. Each grid represents a 50 kbp × 50 kbp square. Regions highlighted with red and blue horizontal bars indicate repeat regions described in the text.

**Figure S3**

Cumulative plots of the number of repeat units detected in raw long reads mapped to (A) 18,816−35,332 bp and (B) 317,602−319,352 bp. Only long reads longer than each region were analyzed.

**Figure S4**

Assembly qualities were compared at different coverages with or without polishing steps. The number of gaps (left) and the number of precisely predicted proteins (right) are shown for (A) assembly by Flye and (B) those polished by Medaka. The definition of a precisely predicted protein is described in Materials and Methods. The high-accuracy base-call mode alone is shown. Each dot represents different subsampling. Red lines indicate the number of predicted genes in the reference MsV T19 genome.

**Figure S5**

Aligned proportion and sequence identity between genomes of newly isolated viruses assembled by different software programs without polishing with short reads. Percentages indicate the fraction of the reference genome shown in the X-axis aligned by the query genome shown in the Y-axis (left panels) or the sequence identity of the query genome shown in the Y-axis to the reference genome shown in the X-axis (right panels).

**Figure S6**

Aligned proportion and sequence identity between genomes of newly isolated viruses assembled by different software programs with polishing by short reads.

**Figure S7**

(A) Total length and (B) N50 of each assembly at different coverages using Flye, Raven, and Miniasm for newly isolated viruses. Dots and colors indicate subsampling and software, respectively. Red labels show the virus isolate names.

**Figure S8**

Heatmap of the reference genomes and newly isolated viruses clustered by ANI values. ANI was calculated by FastANI, and the values below 75% were set as 0%. Accession numbers or isolate names, families and linages of the viruses are shown on the right side of the heatmap. Red names indicate viruses isolated in this study.

**Figure S9**

Phylogenetic trees of (A) BST12E_145 and (B) BST12E_501, constructed with the LG+I+G4 and Blosum62+I+G4 models, respectively. Bootstrap values above 80 are shown. The genes encoded by the pithovirus BST12E are designated by asterisks.

**Table S1**

List of genomes used in this study.

**Table S2**

Summary of the sequencing output.

**Table S3**

Sample sources and tentative names of the newly isolated viruses.

**Table S4**

Summary of *de novo* assembly.

**Table S5**

Results of BLASTX search for short contigs in pithovirus BST12E assembled by Miniasm and Raven.

**Table S6**

Count and proportion of proteins in hybrid assemblies that were predicted in long-read-only assemblies.

**Table S7**

Giant virus genomes showing the highest ANI with assembly of newly isolated viruses polished by short reads.

**Table S8**

ANI between the isolated pithoviruses.

**Table S9**

Annotations of genes exclusively found in BST12E.

## Notes

### Competing Interest Statement

The authors have declared no competing interest.

