## Supplementary figures and images for "A rapid genome-wide analysis of isolated giant viruses only using MinION sequencing"

### Fig. S1

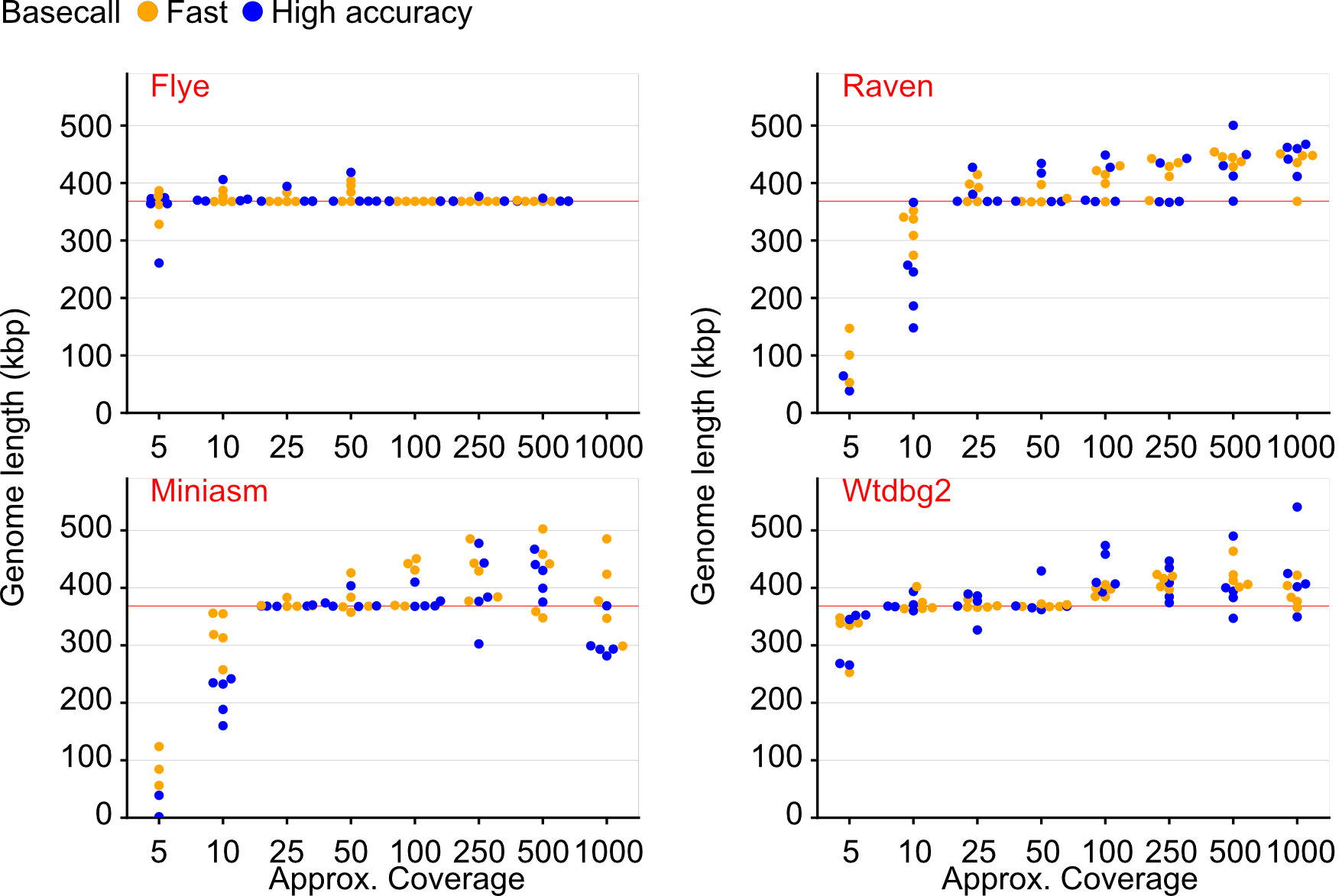

### Fig. S2

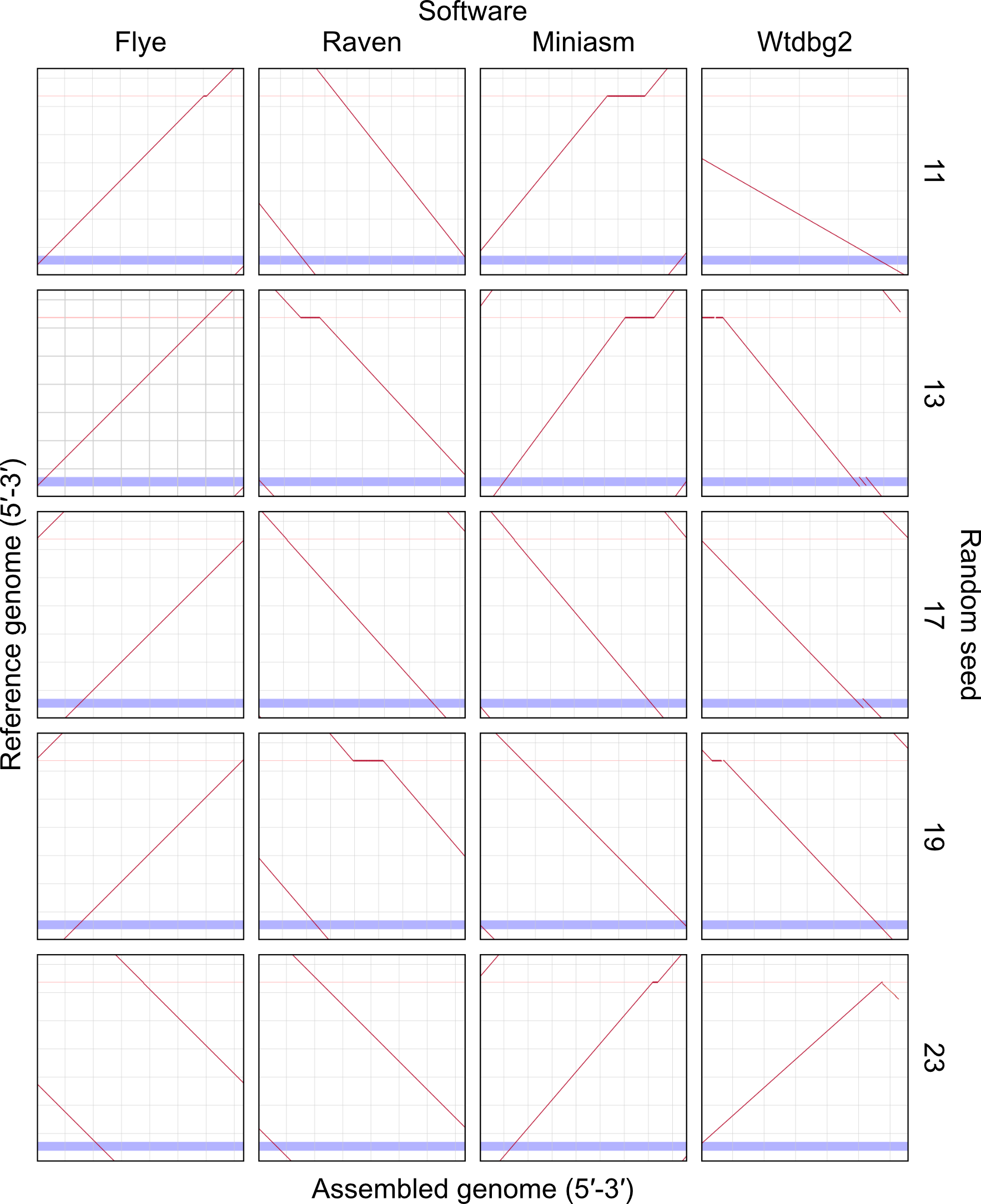

### Fig. S3

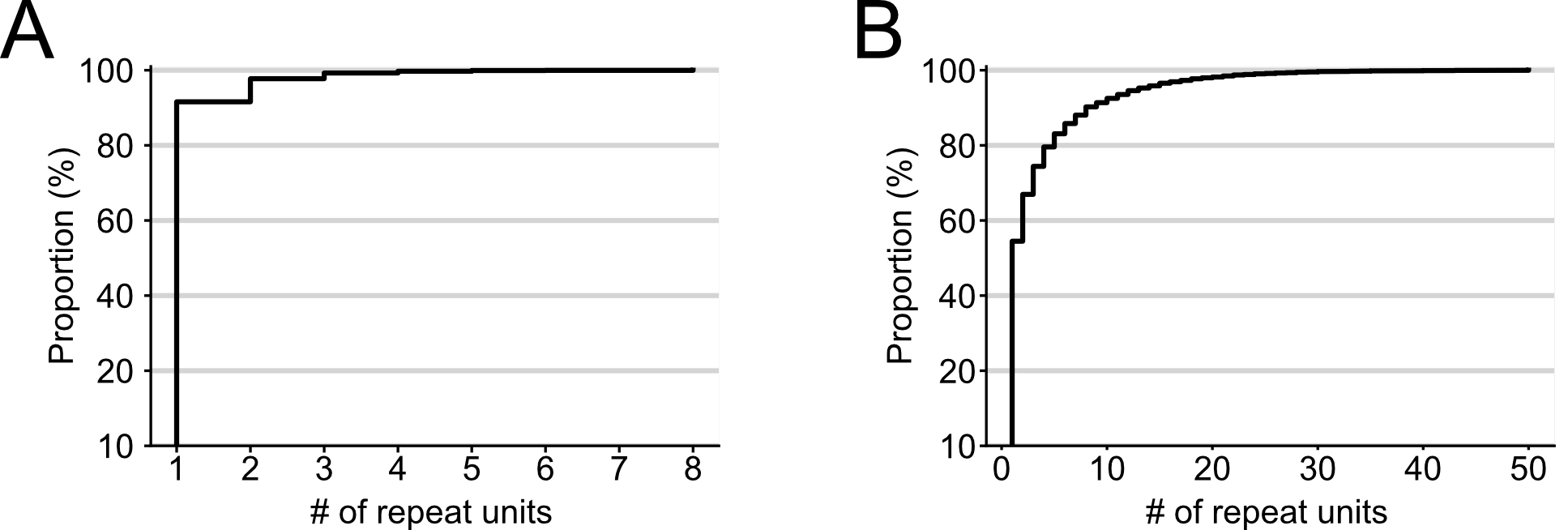

### Fig. S4

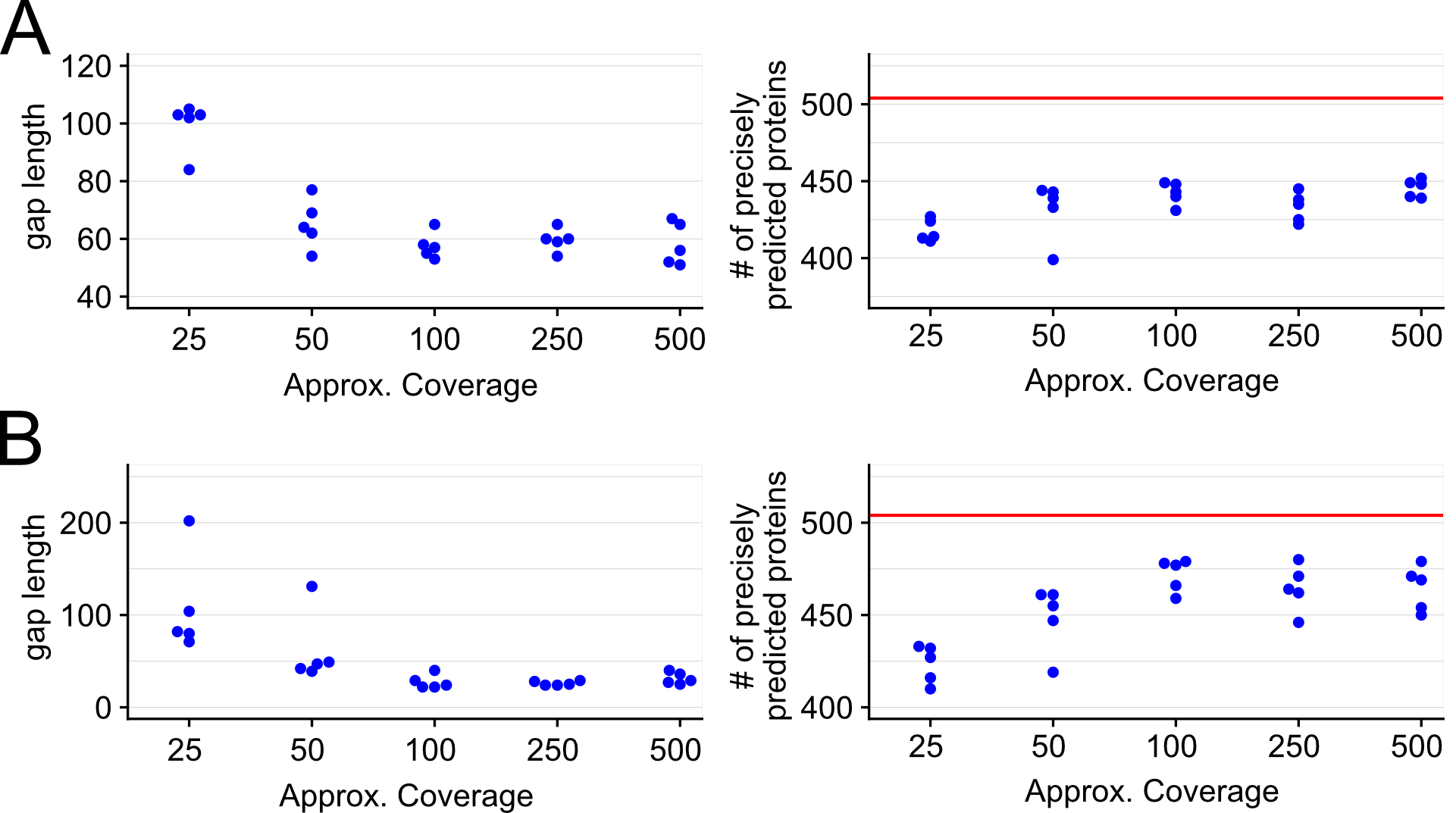

### Fig. S5

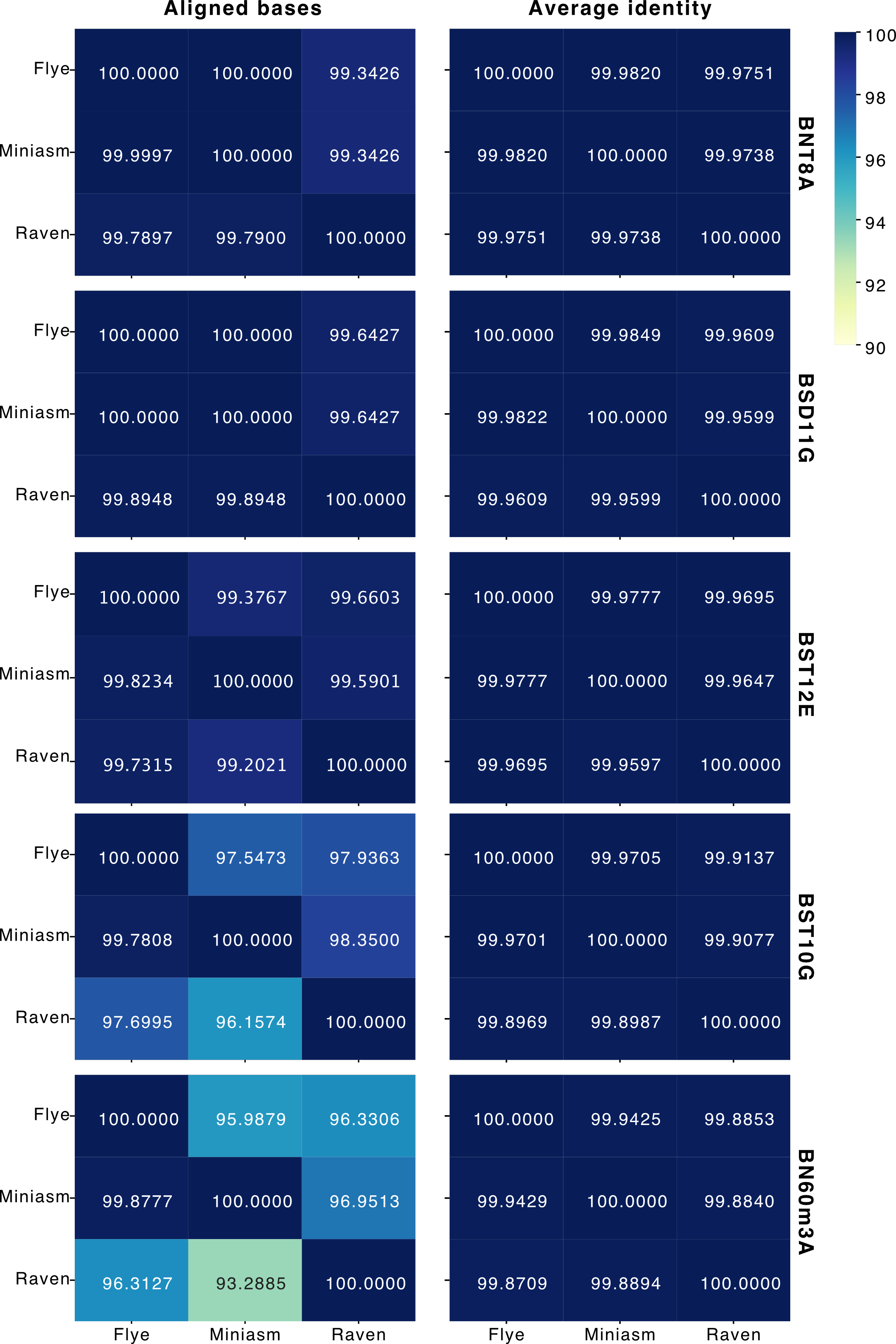

### Fig. S6

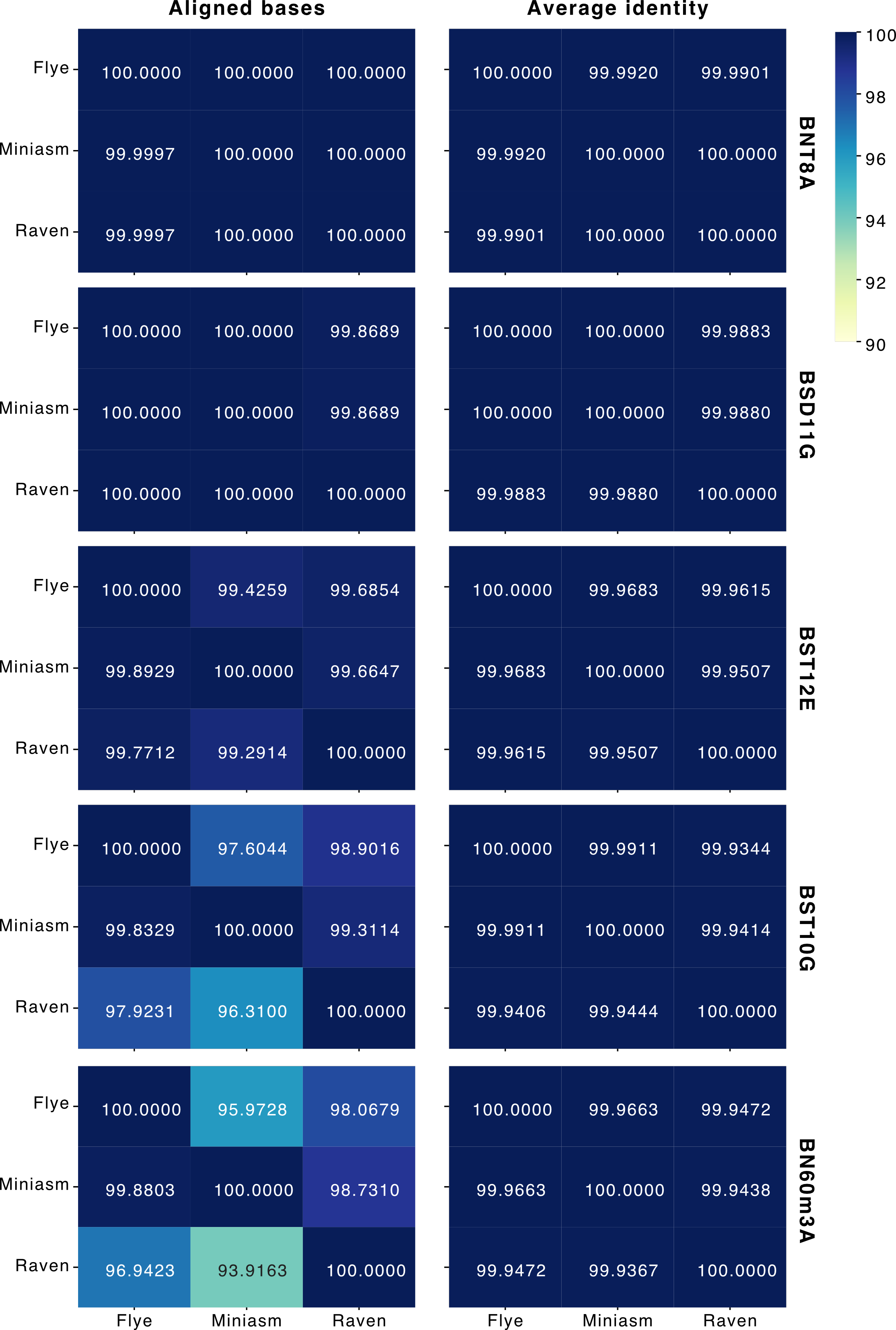

### Fig. S7

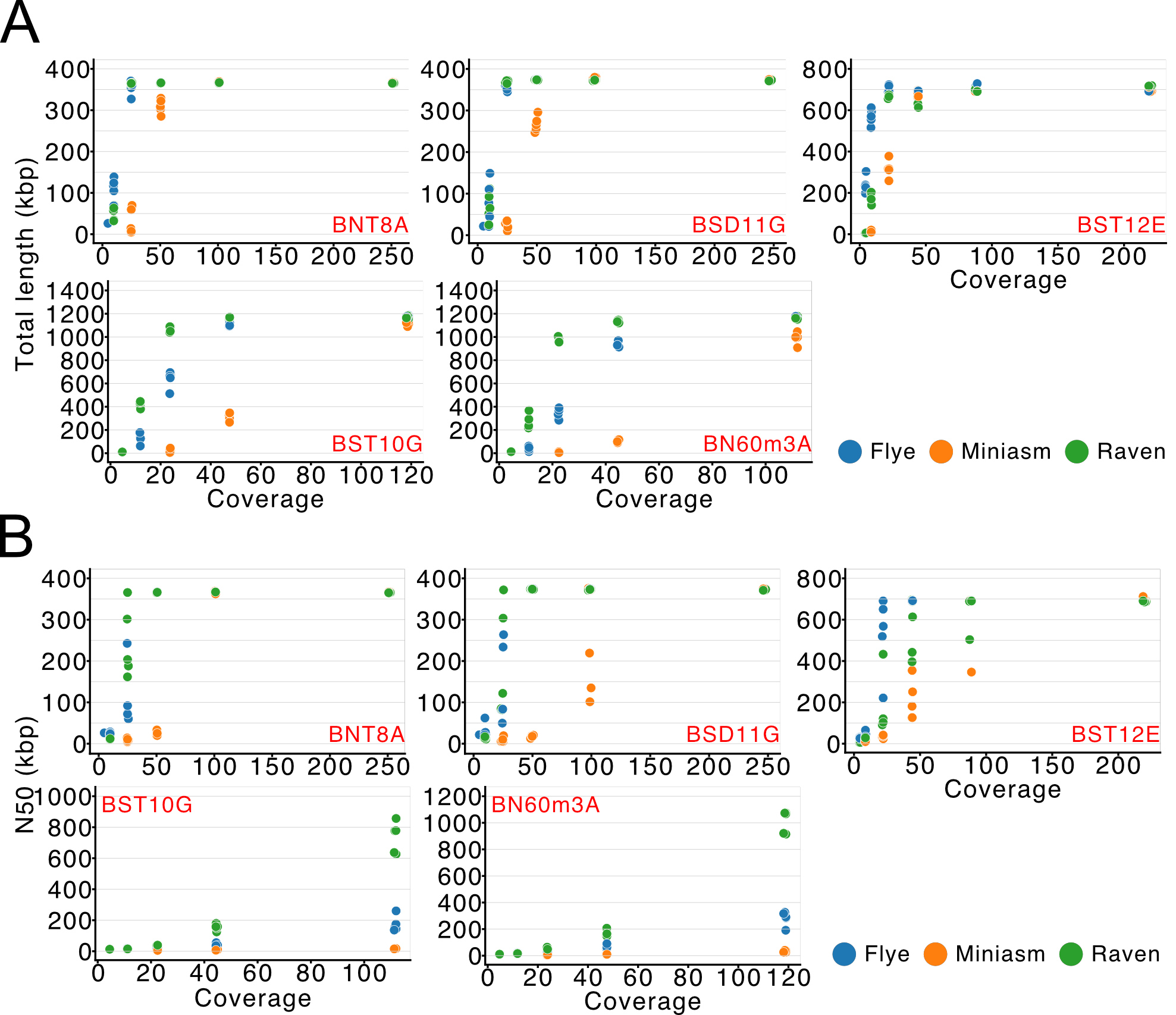

### Fig. S8

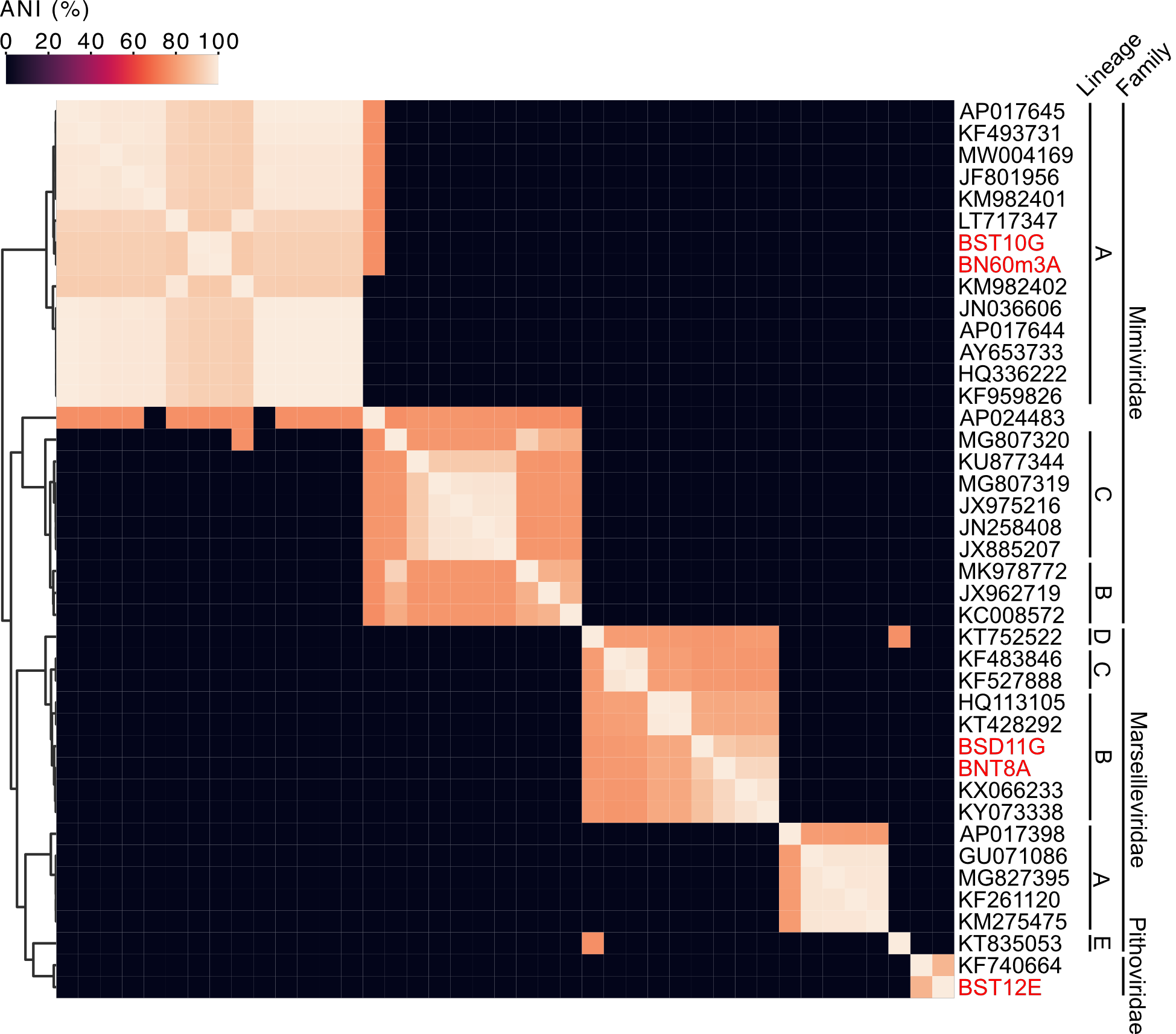

### Fig. S9

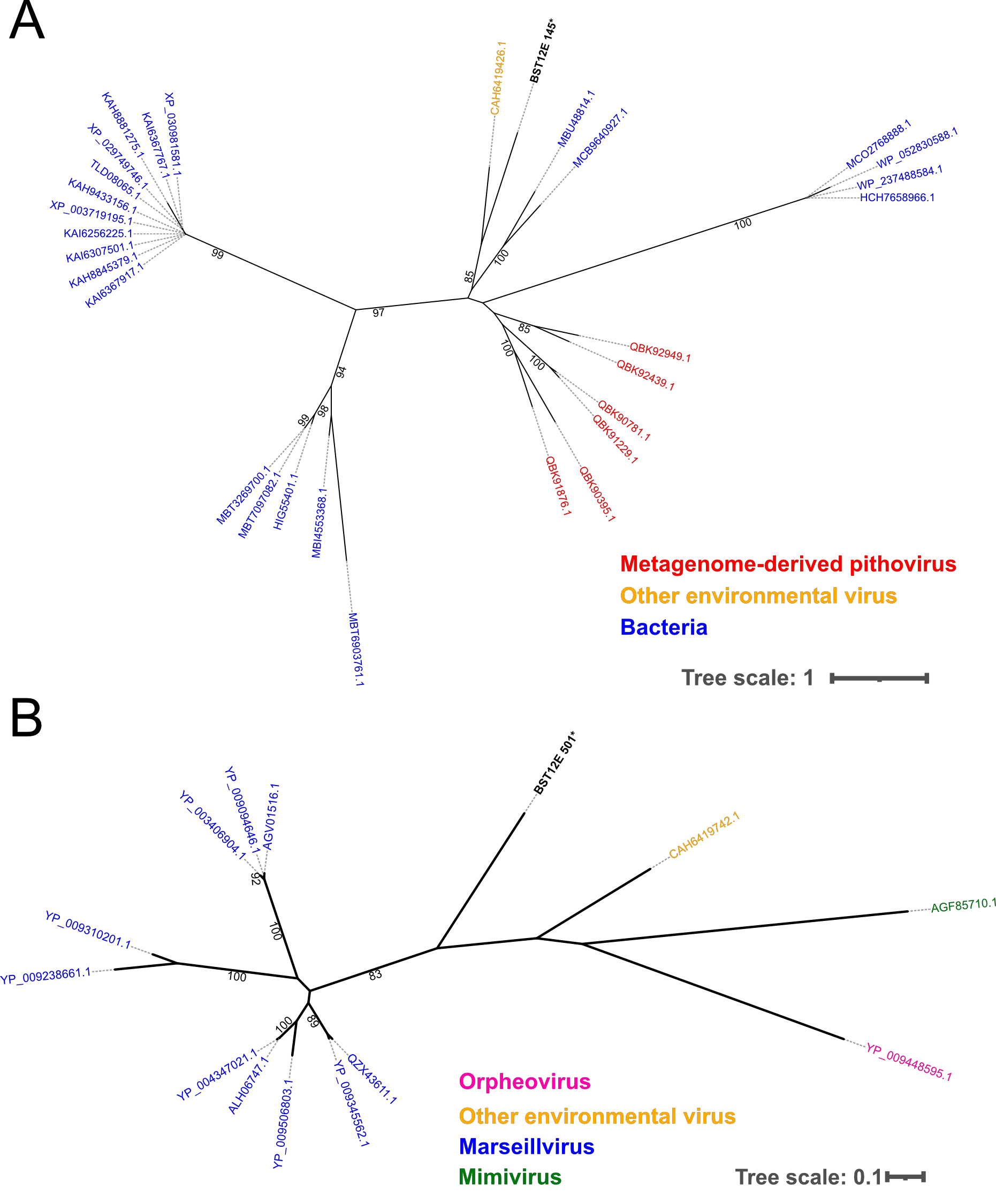

### Table S1

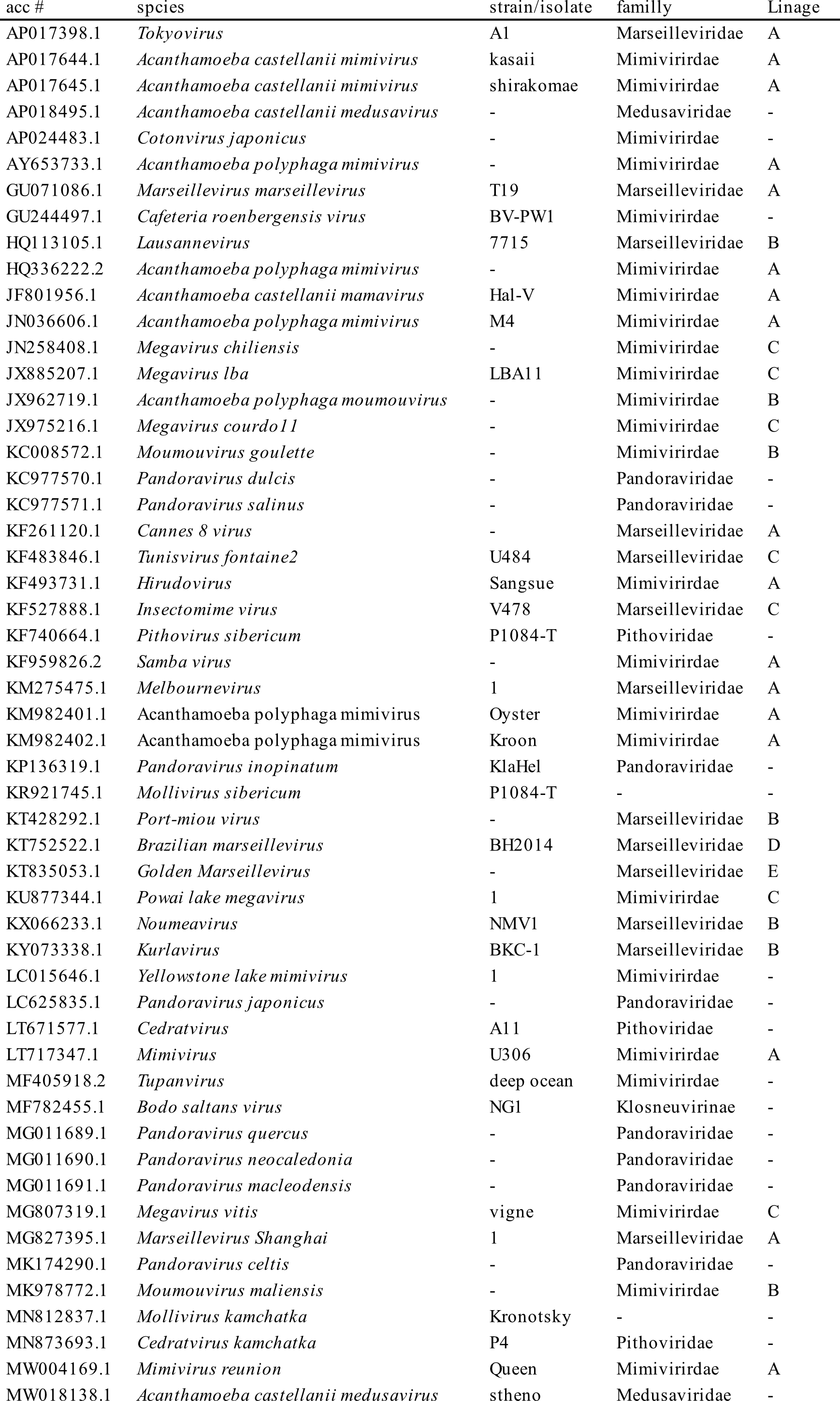

### Table S2

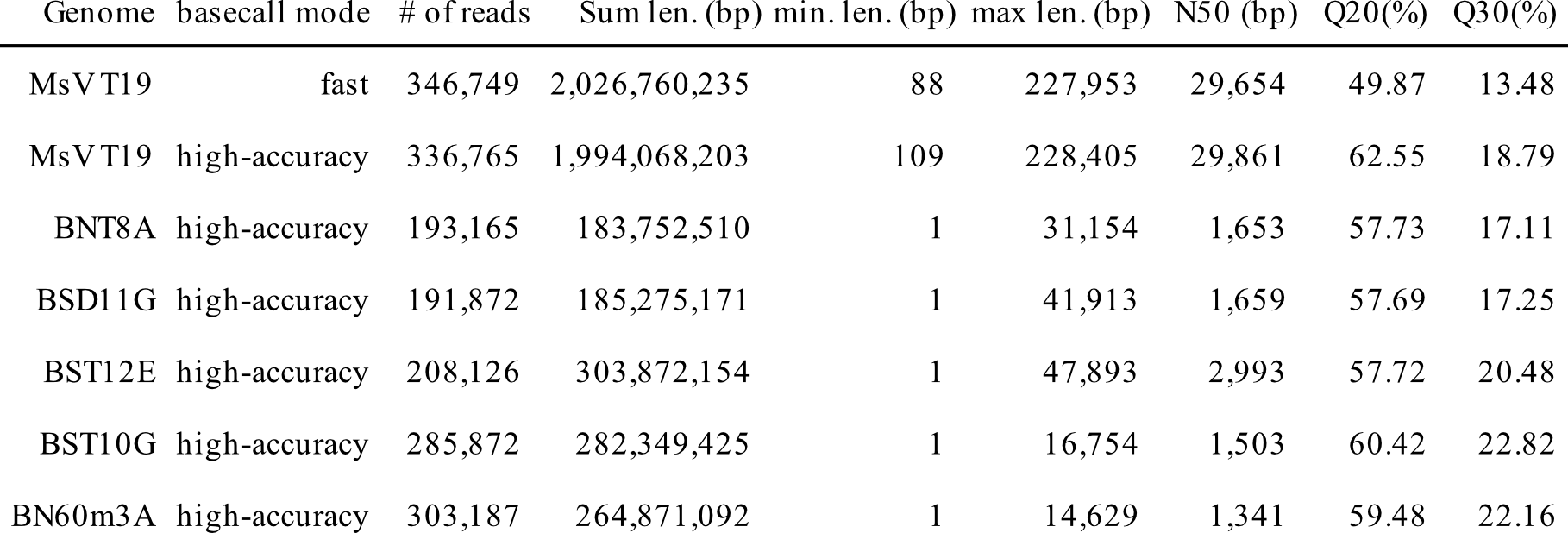

### Table S3

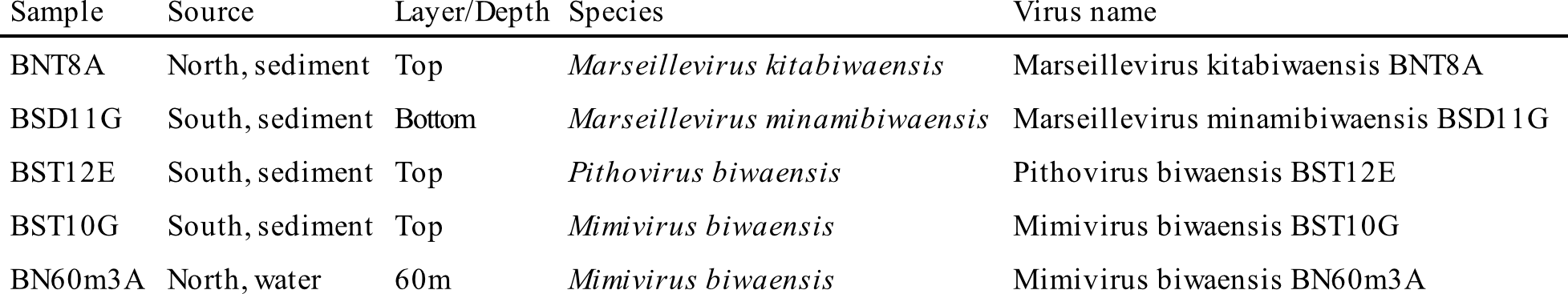

### Table S4

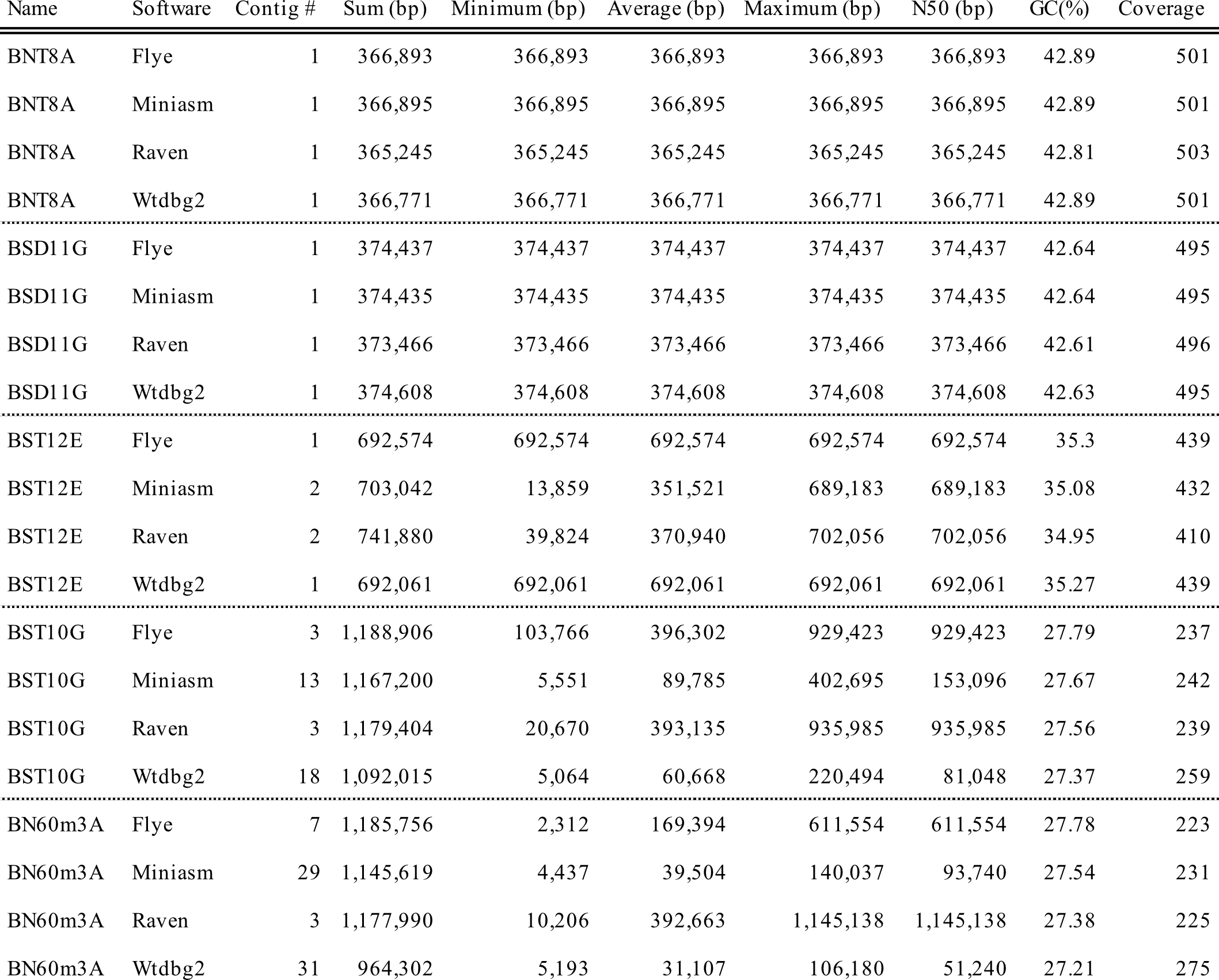

### Table S5

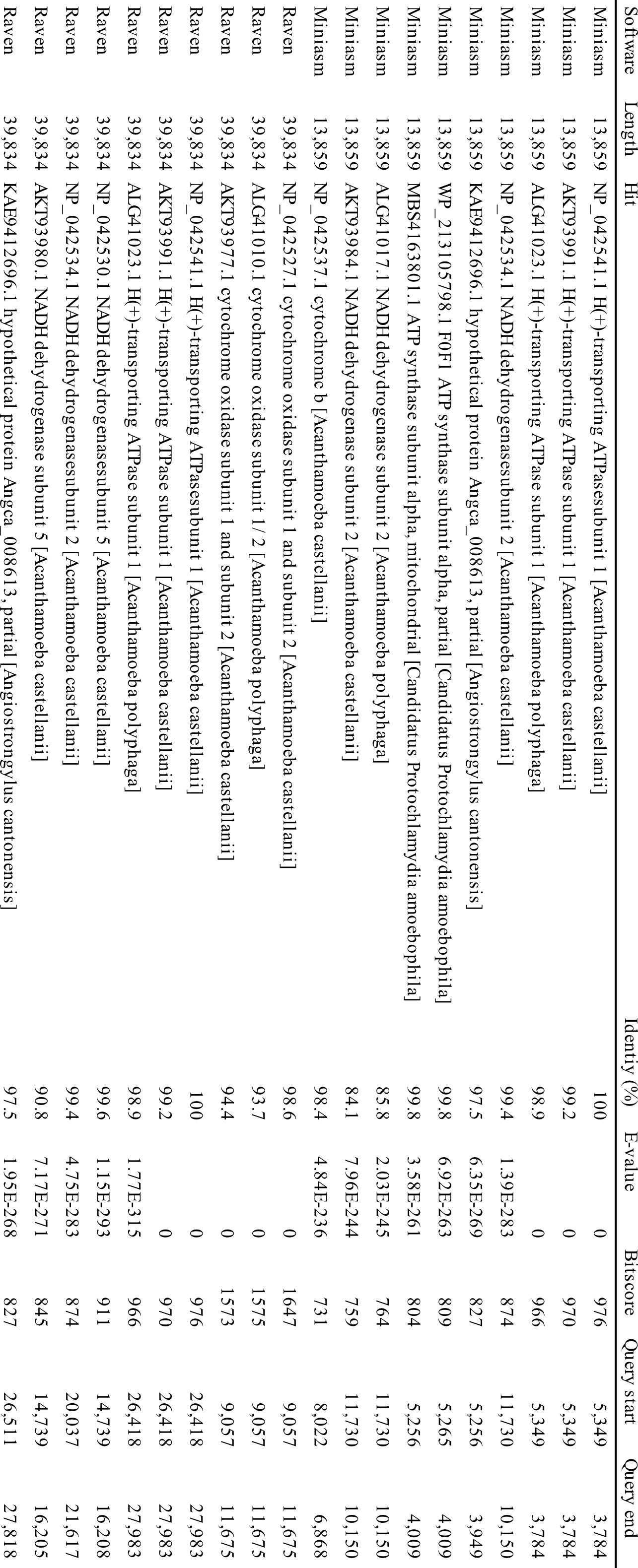

### Table S6

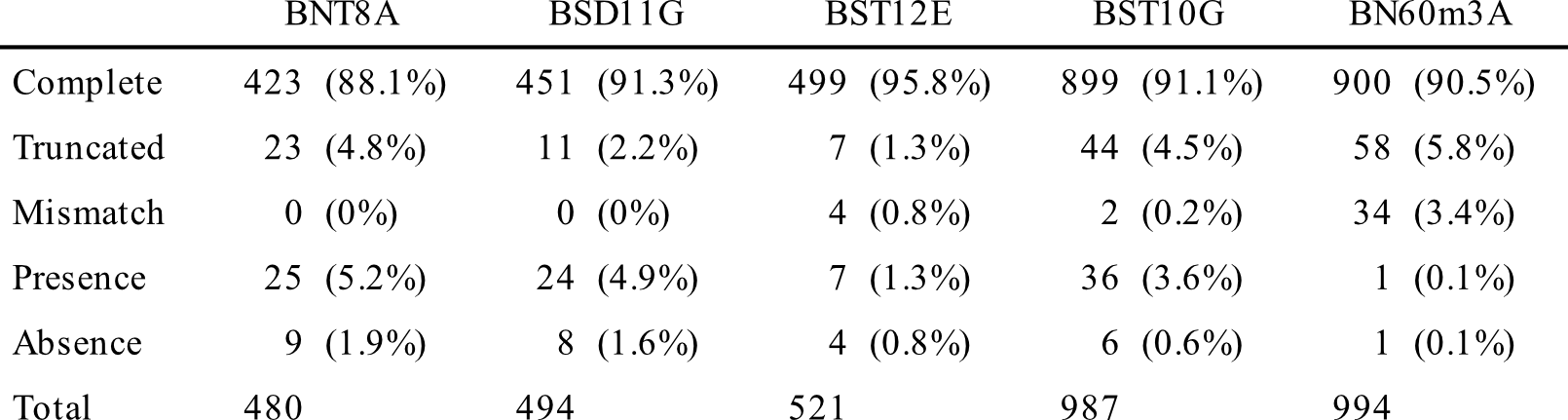

### Table S7

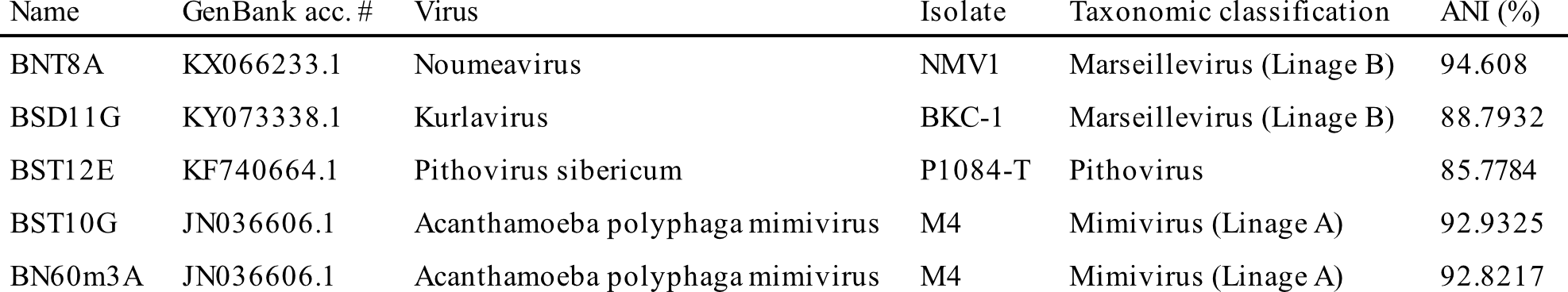

### Table S8

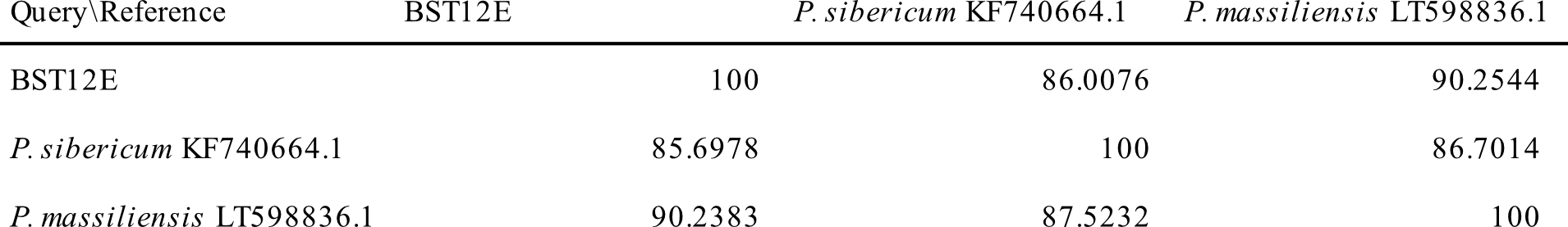

### Table S9

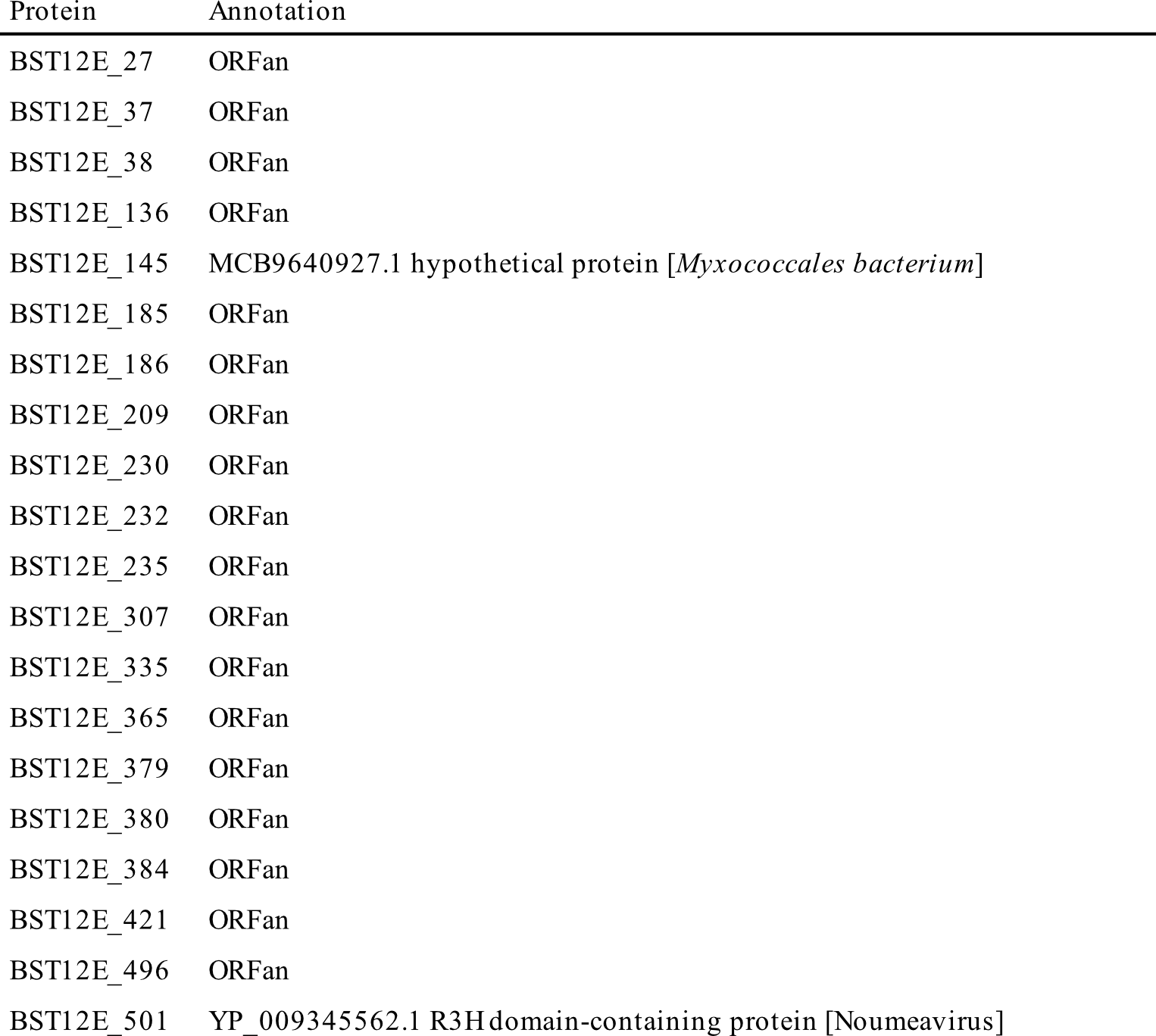
